# Nutrient Signaling-Dependent Quaternary Structure Remodeling Drives the Catalytic Activation of metazoan PASK

**DOI:** 10.1101/2024.06.28.599394

**Authors:** Sajina Dhungel, Michael Xiao, Rajaian Pushpabai Rajesh, Chintan Kikani

**Author notes:** Current address: Cornell/Rockefeller/Sloan Kettering Tri-Institutional MD-PhD Program, New York 10031, USA.

## Abstract

The Per-Arnt-Sim (PAS) domains are characterized by diverse sequences and feature tandemly arranged PAS and PAS-associated C-terminal (PAC) motifs that fold seamlessly to generate the metabolite-sensing PAS domain. Here, using evolutionary scale sequence, domain mapping, and deep learning-based protein structure analysis, we deconstructed the sequence-structure relationship to unearth a novel example of signal-regulated assembly of PAS and PAC subdomains in metazoan PAS domain-regulated kinase (PASK). By comparing protein sequence, domain architecture, and computational protein models between fish, bird, and mammalian PASK orthologs, we propose the existence of previously unrecognized third PAS domain of PASK (PAS-C) formed through long-range intramolecular interactions between the N-terminal PAS fold and the C-terminal PAC fold. We experimentally validated this novel structural design using residue-level cross-linking assays and showed that the PAS-C domain assembly is nutrient-responsive. Furthermore, by combining structural phylogeny approaches with residue-level cross-linking, we revealed that the PAS-C domain assembly links nutrient sensing with quaternary structure reorganization in PASK, stabilizing the kinase catalytic core of PASK. Thus, PAS-C domain assembly likely integrates environmental signals, thereby relaying sensory information for catalytic control of the PASK kinase domain. In conclusion, we theorize that during their horizontal transfer from bacteria to multicellular organisms, PAS domains gained the capacity to integrate environmental signals through dynamic modulation of PAS and PAC motif interaction, adding a new regulatory layer suited for multicellular systems. We propose that metazoan PAS domains are likely to be more dynamic in integrating sensory information than previously considered, and their structural assembly could be targeted by regulatory signals and exploited to develop therapeutic strategies.

---

Period (Per)-Aryl Nuclear Translocator (Arnt)-Single Minded (SIM) (PAS) domains are ancient protein modules found in numerous signaling proteins of physiological and clinical significance across the Tree of Life (ToL) [1–4]. Structurally, PAS domains comprise approximately 60 amino acid-long PAS folds (also termed S1 box) connected to 40-50 amino acid-long PAS-associated C-terminal (PAC) folds (S2 box) via a variable-length linker sequence [1, 5, 6]. Proteins containing PAS domains facilitate diverse signaling and regulatory functions by linking environmental cues to their functional domains [1, 4].

In humans, PAS domain-containing proteins play critical roles in various physiological processes, including circadian rhythm regulation, oxygen sensing, cellular metabolism, and energy balance [3, 4]. PAS domains are encountered in diverse functional contexts and serve as transcription factors such as Period 1-3, Clock, Brain and Muscle ARNT-like 1-4 (Bmal1-4), neuronal PAS domain protein 1-4 (NPAS1-4), Hypoxia Inducible Factor 1α (HIF1α), HIF1β, and Nuclear hormone receptor CoRrepressor 1-2 (NcoR1-2), as well as components of potassium ion channels (KCN 1-8), and serine/threonine kinases such as PAS domain kinase (PASK) [3].

PASK is the only known mammalian protein kinase that combines a nutrient-sensing PAS domain with a serine/threonine-kinase domain. PASK is vital for stem cell differentiation and regulation of organismal metabolism [7–12]. We previously elucidated the crystal structure of the kinase domain of PASK and revealed its activation loop-phosphorylation independent mode of substrate catalysis [13]. Additionally, the solution structure of the PAS-A domain of PASK has been determined, revealing its metabolite-binding capacity [14]. Although PASK exhibits modest activity under unstimulated conditions, nutrient signaling enhances PASK catalytic activity via mechanistic Target of Rapamycin (mTOR) complex 1 (mTORC1)-mediated multisite phosphorylation [11]. Notably, mTORC1 phosphorylates the unstructured region between the PAS-A and kinase domains. However, the structural link between mTORC1-induced phosphorylation and catalytic activation of PASK remains unclear. We hypothesized that mTORC1-stimulated phosphorylation affects the overall 3D structure of PASK, leading to catalytic activation.

Here, by integrating Tree Of Life (TOL)-guided sequence analysis with deep-learning-based structural models, we describe and experimentally validate the nutrient-sensitive assembly of two independent polypeptide chains, PAS and PAC, that together form the third PAS domain (PAS-C) in the metazoan PASK. PAS-C domain assembly is critical for the catalytically stable quaternary conformation of PASK and is targeted by nutrient signaling via mTORC1-mediated phosphorylation. Thus, our findings suggest a transition from a partially unfolded state of the PAS-C domain to a more robust tertiary structure, culminating in the formation of the nutrient-sensitive quaternary structure of PASK that stabilizes the kinase catalytic core. Our analysis provides a unique example of PAS domain formation through long-range intramolecular assembly and paves the way for exploring signal-regulated tertiary structure assemblies in other mammalian PAS domain-containing proteins.

## Results

### Phylogenetic analysis of PASK across metazoans revealed diverse PAS domain distribution patterns

The primary sequences of human and mouse PASK contain one verified PAS domain (PAS-A, aa 132-232) and one serine/threonine kinase domain (kinase, aa 999-1251) [5, 13]. Additionally, the InterPro database [15], which integrates multiple domain signatures from the Conserved Domain Database (CDD) [16] and SMART database [17], predicts the presence of a shorter PAS-B domain (aa 335-402) at a medium stringency level of detection (Figure 1A). The remaining portion of the human PASK sequence (aa 403-998), approximately 600 amino acids, is considered unstructured but functionally significant. This assertion is based on our recent studies, which identified critical regulatory elements within this unstructured region that regulate PASK subcellular distribution, interactions with the substrate Wdr5, and catalytic activation upon mTORC1-mediated phosphorylation [10–12]. Because these biochemical signals primarily target sequences outside the functional domains of PAS-A (PDB-ID: 1LL8) and serine/threonine kinase (PDB-ID: 3DLS), we aimed to understand how these regulatory inputs in the unstructured region of PASK affect the known structural elements that control PASK functions. We hypothesized that functionally important domains and regulatory motifs in unstructured regions ought to be conserved across mammalian PASK.

**Figure 1.**
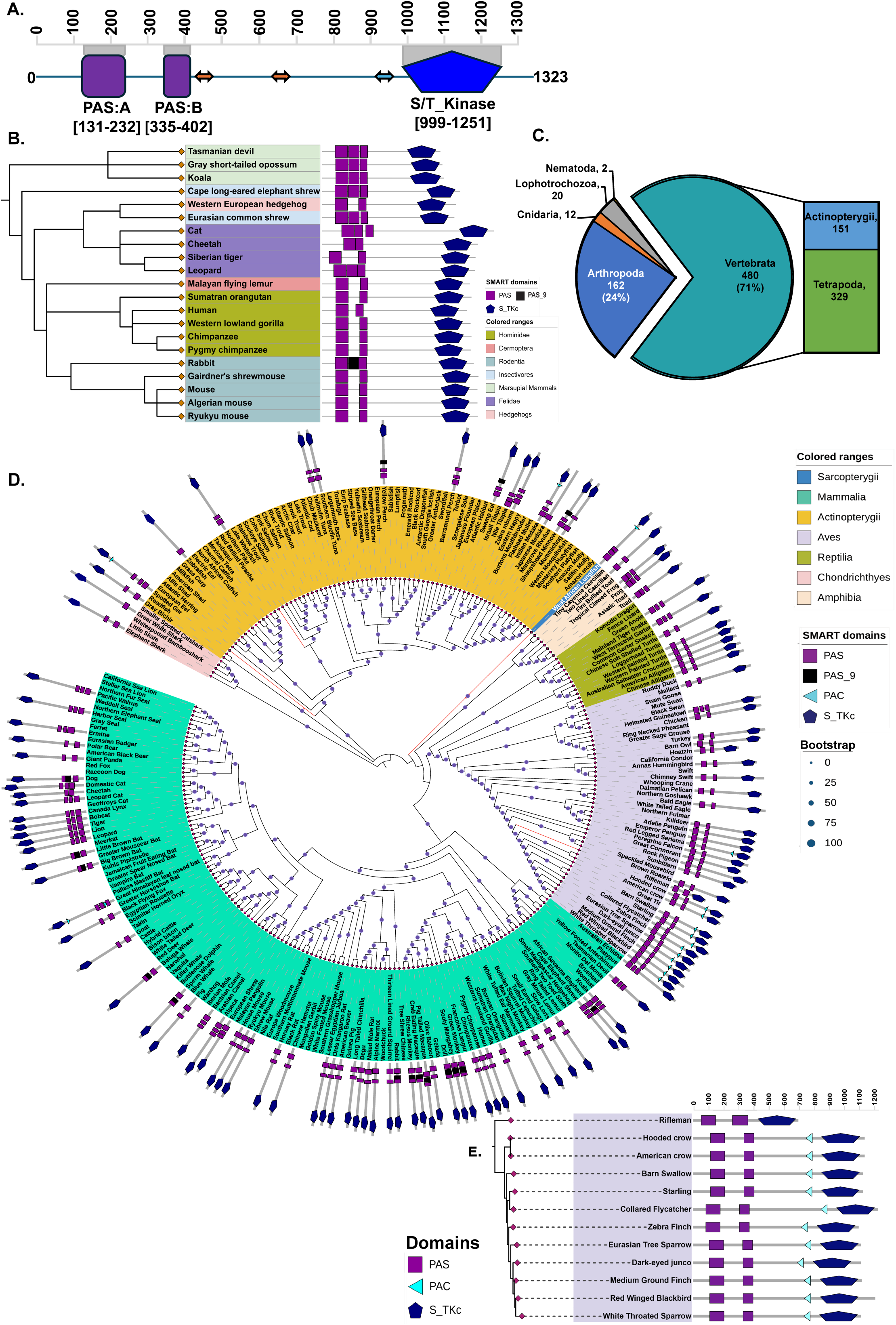
Domain mapping across the tree of life revealed the diversity of the domain architecture in PASK. **A.** Domain architecture of human PASK as predicted by the InterPro, SMART, and CDD databases. **B.** Domain architecture of selected mammalian PASK from the SMART database. **C.** Distribution of PASK across the metazoan tree of life. **D.** The maximum-likelihood tree for amino acid sequences inferred from complete PASK sequences for jawed vertebrates was constructed using the JTT+F+R5 (BIC approximation) model in IQ-Tree2. The circle size at each node represents the bootstrap values calculated with 1,000 replications. Domain mapping for the selected jawed vertebrate species was performed using SMART. **E.** Maximum-likelihood tree of PASK from Passeriformes, with the dashed line representing branch length.

To explore this hypothesis, we aligned available mammalian PASK and performed domain-mapping analyses. Most mammalian PASK sequences exhibit a high degree of sequence conservation and domain distribution patterns similar to human PASK, with the PAS-A domain being approximately 100 amino acids long and the putative PAS-B domain being approximately 60 amino acids long. However, combined domain mapping using SMART, InterPro, CDD, and HMMSEARCH identified an additional PAS domain in several mammalian PASK sequences belonging to either the canonical PAS or PAS_9 family located between the PAS-A and putative PAS-B domains of human PASK (Figure 1B, Figure S1). Across all mammalian species examined, the domain corresponding to the human PAS-B domain (depicted in Figure 1A) is shorter, consisting of approximately 60 amino acids, compared to the typical length of approximately 100 residues found in the canonical PAS domain, regardless of whether it is configured with two or three PAS domains.

PAS domains are characterized by diverse sets of sequence signatures, which could lead to an under-representation of PAS domains within the proteome when examined using sequence-based domain-mapping algorithms. We theorized that examining domain architecture across the Tree of Life (TOL) could uncover new insights into the distribution, structure, and evolution of PASK in general, and for shorter PAS-B domains in particular. To this end, we performed phylogenetic analysis of PASK across metazoans. Orthologs of PASK have been identified throughout the eukaryotic tree of life, with approximately 700 PASK genes identified in the metazoans of approximately 650 species for which transcriptome or genomic data are available. As 71% of all metazoan PASK genes were found in chordates (Figure 1C), we focused our analysis on diverse taxa of jawed vertebrates from publicly available genome/transcriptome databases covering cartilaginous and bony fish, amphibians, birds, and mammals. We aligned the PASK sequences from jawed vertebrates and generated a maximum-likelihood tree. Our analysis revealed that PASK is present in sharks, ray-finned- and lobed-fin fish, lungfish, amphibians, reptiles, birds, and mammals. It is present in evolutionarily important species such as the spotted gar (Lepisosteus oculatus), West African lungfish (Protopterus annectens), northern pike (Esox lucius), and bird rifleman (Acanthisitta chloris). We then performed a domain analysis for selected species across broad taxonomic clades using HMMER-based SMART, combining the predicted domain distribution with a phylogenetic tree (Figure 1D). Similar to mammalian PASK, domain mapping algorithms have identified one, two, or three PAS domains and a single C-terminal serine/threonine kinase domain for most jawed vertebrates. Consistent with mammalian PASK, the PAS-A domain of PASK in the jawed vertebrate species was predicted to be approximately 100 amino acids in length. In contrast, the PAS-B domain in the analyzed species was approximately 60 amino acids in length, except for rifleman, whose PAS-B domain was also approximately 100 amino acids long. In addition to these two domains, several species of jawed vertebrates also contain a PAS domain from the PAS_9 family between the PAS-A and PAS-B domains, according to SMART prediction.

In addition to these PAS domains, SMART also identified a single PAS-associated C-terminal (PAC) motif-like sequence in PASK from approximately 16 jawed vertebrate species (Figure 1D). Where predicted, a single PAC motif was found at a consistent location, approximately 63-69 amino acids upstream of the kinase domain. An exception was found in the Himalayan leaf-nosed bat (*Hipposideros armiger*), for which SMART predicted two PAC motifs: the N-terminal PAC motif split from the PAS-A domain and the other C-terminal PAC motif upstream of the kinase domain (Figure 1D, cyan triangles).

PAC motifs are integral to the PAS domain architecture. It is generally accepted that PAS domains consist of two distinct subdomains: PAS-fold (or S1 box) and PAC-motif (or S2 box) [5, 6], which fold together to form the canonical PAS domain, as illustrated in Figure S2. PAS-fold is approximately 60 amino acids in length and comprises of two antiparallel beta-sheets (labeled Aβ and Bβ; depicted in purple in Figure S2) and four alpha-helices (Cα, Dα, Eα, and Fα; depicted in purple in Figure S2). The PAS fold is connected to the PAC motif via a variable-length linker loop region. The PAC-motif is approximately 40 residues long and contributes three antiparallel beta-sheets Gβ, Hβ, and Iβ (shown in cyan in Figure S2) to the PAS domain architecture. The PAC fold is centrally located, houses a ligand-binding pocket, and plays a role in the allosteric functions of the bacterial PAS domain (Figure S2). Although typically annotated as separate entities in most protein domain annotation programs, the PAS fold and PAC motifs are arranged in tandem; consequently, they are redefined as a single cohesive folding unit, resulting in canonical PAS domains comprising approximately 100 residues [18].

In proteins of human origin that contain PAS domains, such as bHLH transcription factors, domain prediction tools such as InterPro and SMART (based on HMMER) have consistently found tandem PAS-PAC folds in the second (PAS-B) domain, which folds together to span a typical length of 100 amino acids, as shown in Figure S3A-B. However, unlike bHLH transcription factors, neither SMART nor InterPro identified any PAC-like sequences next to the PAS-B domain in any metazoan PASK (Figure 1D, S3A).

In certain species where SMART did identify a PAC-like sequence in PASK (Figure 1D-E), this sequence was separated from the PAS-B domain by a long, unstructured loop. For these PAC domains, we found no evidence of any immediate N-terminal sequences adjacent to them that could fold with the PAC motif to form the standard PAS domain. Moreover, we did not observe any specific evolutionary pattern in the prediction of PAC-like motifs across metazoan PASK, which includes both invertebrates and vertebrates (Figure S4). The only exception was passerine birds, for which SMART predicted a PAC domain in almost all representative members of our PASK sequence database, separated from the shorter PAS-B domain (Figure 1E). Interestingly, rifleman, a common ancestor of passerine birds, did not contain a predictable PAC-like motif in its PASK sequence (Figure 1E). However, its PAS-B domain length matches that of canonical PAS domains (∼100 amino acids) compared to other passerine birds and most metazoan PASK (∼60 residues) (Figure 1E). Thus, phylogeny-guided domain analysis suggests a significant diversity in the number and length of PAS domains across vertebrate PASK.

### Structural phylogenetic analyses reveal the unusual PAS domain folding and assembly mechanisms

To understand how diverse domain distribution patterns and sequence lengths are reflected in PASK protein structures, we analyzed the available AlphaFold structures of PASK for selected metazoan species. We conducted a comparative structural analysis of PASK across a variety of species, domain patterns, and lengths using AlphaFold 2.0 (AF2). Because the structure of rifleman PASK is not available in the AlphaFold database, we generated its structural model using AlphaFold2 run on CoLabFold [19]. Our structural phylogenetic analysis suggests that regardless of inconsistencies in assigning the number and length of PAS domains by any sequence homology-based algorithm, all metazoans PASK contain three PAS-like domains of approximately 100 residues each and a canonical kinase domain (Figure 2A). Thus, AF2 structures from diverse species, such as spotted gar, rifleman, zebra finch, and human PASK, exhibited the presence of three PAS domains and one serine/threonine kinase domain (Figure 2A). To analyze the quality of the AF2 models, we compared the predicted local distance difference test (pLDDT) matrices of the AF2 structural models for human and rifleman PASK (Figure 2B-E). In human PASK, relatively high pLDDT scores (≥70) corresponded to three well-defined PAS domains, and scores above 90 corresponded to a well-conserved kinase domain (Figure 2B-C). Within each PAS domain, many residues showed high pLDDT scores, indicating the presence of well-structured regions. Based on comparative sequence-structure analysis in this manuscript, we revised the nomenclature of the PAS domain architecture of PASK across metazoa, which aligns with the SMART-predicted domain boundaries, the PAS-B domain, which is modeled in AF2 structures but is predicted sporadically across metazoan PASK by SMART (PAS_9 family, Figure 1D), and the PAS-C domain, a shorter PAS domain predicted in all metazoan PASK, was previously designated as the PAS-B domain (Figure 1A).

**Figure 2.**
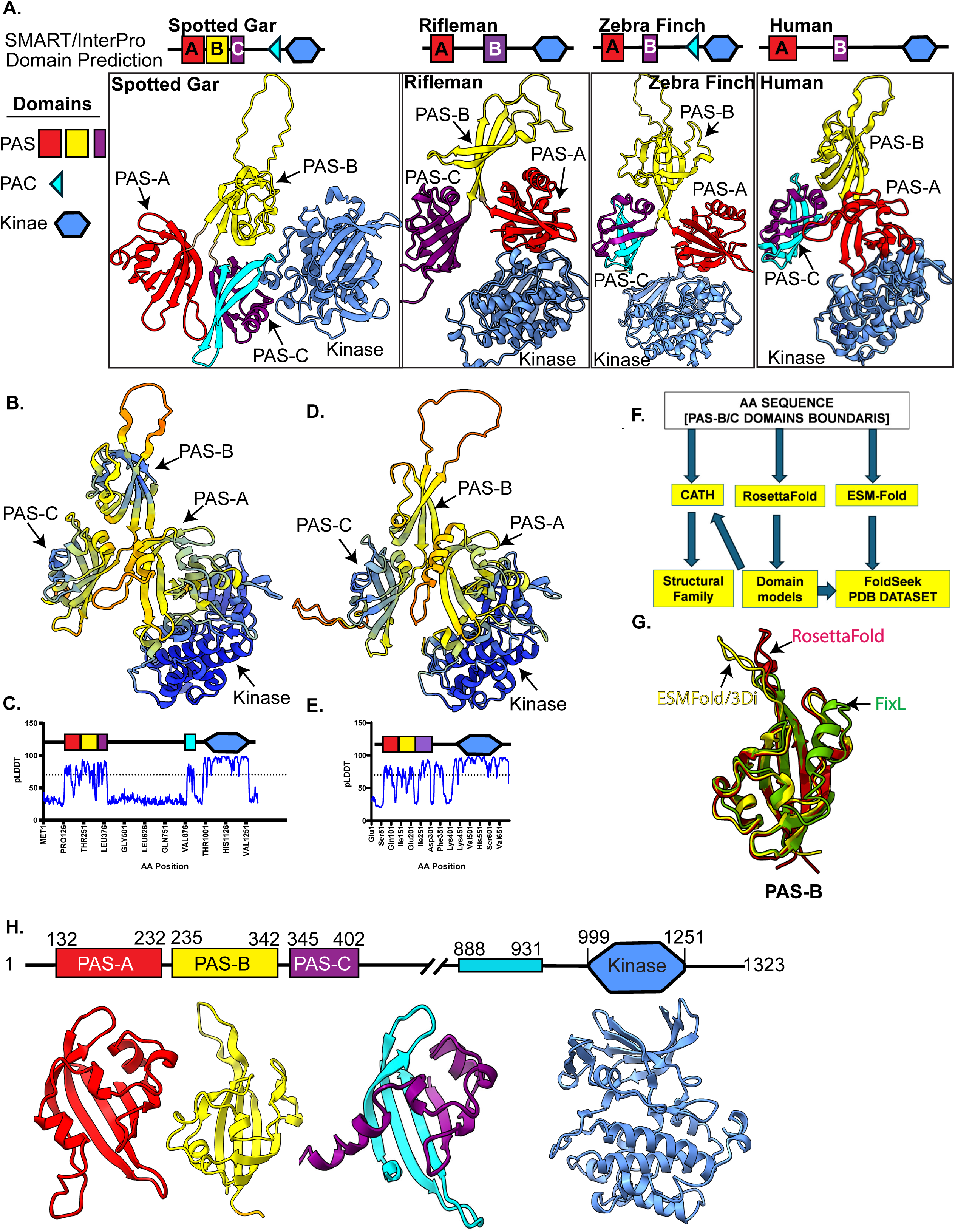
Analysis of Predicted 3-Dimensional Structure of PASK in Jawed Vertebrates. **A.** Domain architecture as predicted by SMART for selected vertebrate PASK (top) compared to their corresponding predicted AlphaFold structures (bottom), highlighting variability in domain prediction versus the commonality of the AlphaFold structural models. **B.** AlphaFold structure of human PASK, colored according to the pLDDT scores and depicting four main protein domains. **C.** Per-residue pLDDT score profiles for human PASK, with pLDDT scores overlaid on the domain boundary map obtained from the AlphaFold structure. **D.** AlphaFold structure of rifleman PASK, colored according to the pLDDT scores and depicting four main protein domains. **E.** Per-residue pLDDT scores profile for rifleman PASK, with pLDDT scores overlaid on the domain boundary map obtained from the AlphaFold structure. **F.** Computational validation workflow to independently validate structural models for PAS-B/C domains and identify the structural family. **G.** Structural overlay of the PAS-B domain model obtained from RosettaFold (Red), ESMFold (Yellow) with the *Bradyrhizobium japonicum* FixL PAS-B domain. **H.** Structure-guided revised domain boundaries of human PASK, noting that two distinct regions in the PAS-C domains contribute to the overall PAS fold.

While the PAS-A and PAS-C domains of rifleman PASK showed good structural models with high pLDDT scores (≥70) (Figure 2D-E), the PAS-B domain did not model well into the structural fold, likely due to missing key residues from the current sequence (Figure S5B). In contrast, the PAS-A and PAS-B domains from selected fish, birds, and mammals exhibited a well-modeled PAS domain architecture and high sequence similarity with human PASK (Figure S5A-B).

Because the solution structure of the PAS-A domain has been well studied [14], we focused on characterizing and validating AF2 models of the PAS-B and PAS-C structures. We developed an independent model validation workflow to compute sequence-guided model analyses to determine whether the predicted PAS-B domain structures conform to a canonical PAS fold (Figure 2F). First, we used a sequence corresponding to the PAS-B domain of human PASK (aa 235-342) to identify the matching Functional Family (FunFamily) in the CATH database. We then performed structural alignment of the human PAS-B sequence in the PDB database using FoldSeek [20]. For this purpose, we directly modeled residues corresponding to the putative PAS-B domain sequence with ESMfold [21] and searched the PDB database in the 3D interaction (3Di) or TM-Align mode without taxonomic restrictions to capture broad, evolutionarily relevant structural homologs. We also generated a RosettaFold [22] structure in template-free mode from a sequence corresponding to the PAS-B domain and analyzed the resulting structure files using either FoldSeek or CATH to identify the structural neighbors of the RosettaFold model. These independent and overlapping analyses produced a CO-bound PAS domain structure from Bradyrhizobium japonicum FixL (PDB ID: 1xj4-A, RMSD: 3.14) and a series of PAS domain structures from hypoxia-inducible factor 1a (PDB ID: 1p97; RMSD: 3.38, 8ck3_A; RMSD: 3.55, 3f1p_A; RMSD: 4.2) as the best scoring models (TM score ≥0.69, probability ≥0.90) for the PAS-B domain (Figure 2G). The AlphaFold structure of the PAS-B domain in CATH and FoldSeek also returned PAS domains from bacterial FixL, human HIF1α, or HIF1β as top hits. These analyses suggest that PAS-A (highlighted in red, aa 132-232 for human PASK) and PAS-B (highlighted in yellow, aa 235-342 for human PASK) are formed from a contiguous sequence that folds into independent canonical secondary structural elements that constitute the PAS domains (Figure 2H). Both domains were approximately 120 amino acids long and contained antiparallel five-stranded β-sheets interspersed with α-helices (Figure 2H).

Compared to the PAS-A and PAS-B structural models, the PAS-C domain exhibited an unusual mode of assembly in nearly all metazoan PASK studied, except for rifleman (Figure 2A). The N-terminal PAS fold contributes approximately 53 amino acids (aa 345-402 for human PASK; purple domain architecture and tertiary structure in Figure 2H). This region matches the smaller PAS domains predicted by CDD and SMART for human PASK (Figure 1A, 2H). The PAS fold sequence is complemented by three antiparallel β-sheet secondary structures contributed by an approximately 43 amino acid stretch between residues 888-931 (Figure 2H, tertiary structure elements in cyan). In contrast to the human and other metazoan PAS-C domains of PASK, the rifleman PAS-C domain is predicted to be as long as the canonical PAS domain (∼100 amino acids) and to exhibit contiguous PAS-C domain folding, which is supported by the AlphaFold structure (Figure 2D-E). Thus, despite the differences in the predicted domain sizes (Figure 1D-E, Figure 2A), a well-modeled PAS-C domain was present in all metazoan PASK domains examined. The unusual yet conserved mode of PAS-C domain assembly between rifleman and other metazoan PASK prompted us to further evaluate and validate the structural model of the PAS-C domain assembly.

### Identification of PAC-motif across metazoan PASK that facilitates PAS-C domain assembly

The PAS-C domain models from humans and rifleman showed remarkable structural congruency despite vastly different domain size predictions and the locations of residues in the primary sequence that generate the tertiary structure (Figures 2B, 2H). To understand the sequence/structure relationship driving the predicted PAS-C domain folding, we compared the PAS-C domains of rifleman and human PASK to identify sequence features contributing to PAS folding in both species (Figure 3A-B). The rifleman PAS-C domain spans amino acids 251-363. The 56 N-terminal residues (aa 251-307) form PAS-fold structures and contribute to Aβ, Bβ, Cα, Dα, Eα, and Fα in the rifleman PAS-C domain (Figure 3A, colored in purple). These residues are also highly conserved in the human SMART/InterPro-predicted PAS-C domain (56 residues, 345-401 aa in humans; Figure 3A-B, colored in purple). The remaining three antiparallel beta-sheets that constitute Gβ, Hβ, and Iβ of the rifleman PAS-C domain are contributed by 43 residues spanning amino acids 320-363 (Figure 3A-B, colored in cyan). These residues showed a high level of sequence identity with the 43 residues between amino acids 888 and 931 of human PASK and contribute to the antiparallel beta-sheets, Gβ, Hβ, and Iβ (Figure 3A-B, colored in cyan). Thus, in human PASK, the N-terminal PAS fold (Figure 3A-B, colored in purple) is separated from the C-terminal PAC fold (Figure 3A-B, colored in cyan) by 486 amino acids and is modeled to fold together to generate a canonical PAS-C domain that is approximately 100 amino acids long (Figure 3B).

**Figure 3.**
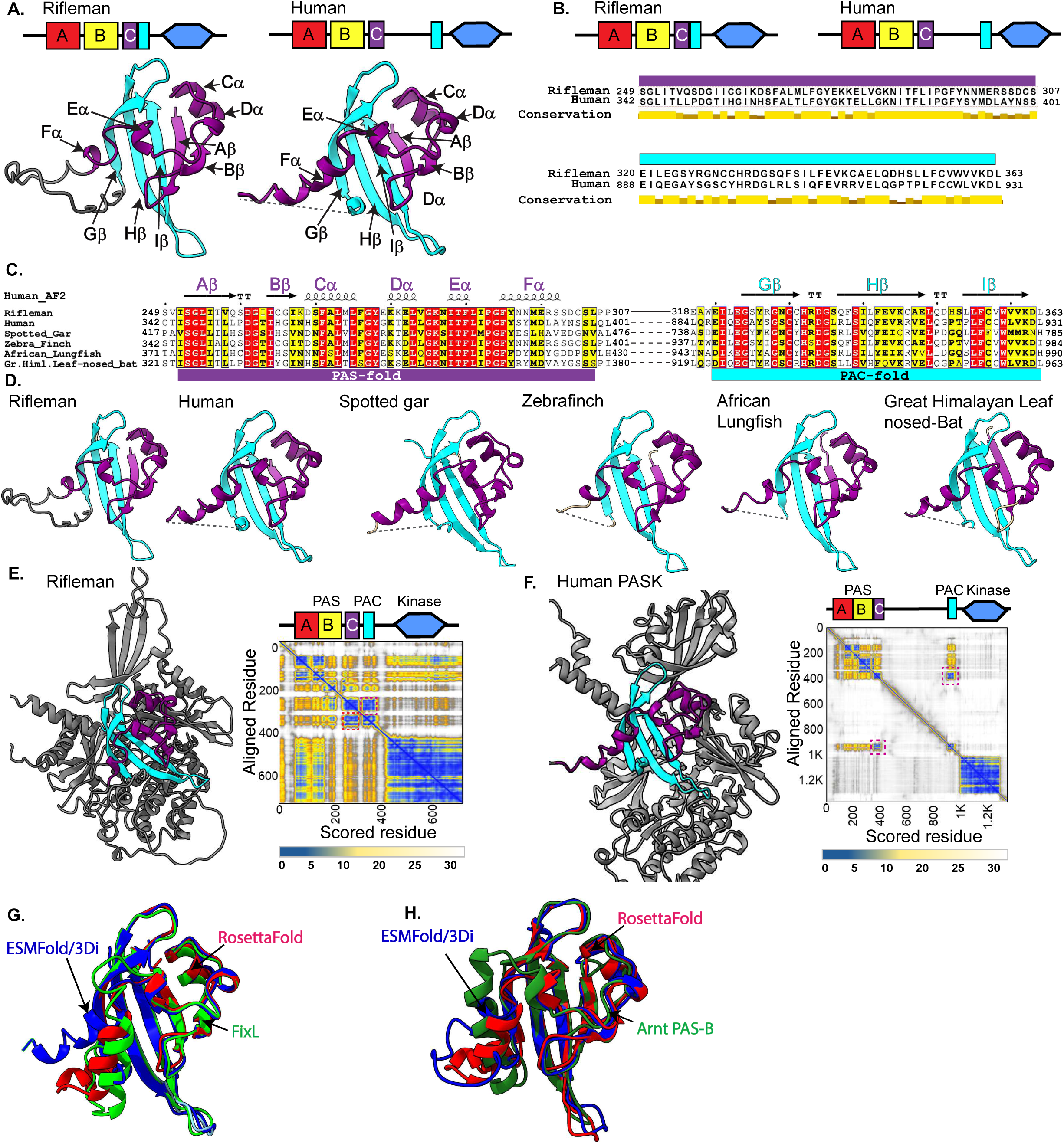
Intramolecular assembly of PAS-PAC folds in the PAS-C domain of PASK. A. Conservation of the structural elements of the PAS-C domain between rifleman and human PASK, with N-terminal PAS-folds in purple and the C-terminal PAC motif depicted in cyan in the structure and domain architecture. **B.** Primary sequence alignment between rifleman and human PASK showing the PAS and PAC fold domain boundaries. **C.** Structure-based alignment of selected vertebrate PASK using the human PASK PAS-C domain as a structural template, with intervening loops of variable length between PAS folds and PAC folds denoted by a dashed line, except for rifleman, which shows a much shorter flexible loop region (denoted by a solid line) connecting the PAS and PAC folds. Sequence alignments were generated using MAFFT and superimposed on the structural elements using EsPript 3.0. **D.** PAS-C domain comparison between representative PASK from fish, birds, and mammals, noting the flexible loop region in rifleman PASK connecting PAS-folds (purple) and PAC-folds (cyan), in contrast to the separation of the PAC-fold from PAS-folds in other species via long intervening loops (dashed line). **E.** Coloring of human PASK structure according to selected residues (magenta square) in the PAE scoring matrix, with dashed squares in the PAE matrix denoting the X and Y positions of residues contributing to PAS-C folds. **F.** Coloring of rifleman PASK structure according to selected residues (magenta square) in the PAE scoring matrix, with dashed squares in the PAE matrix denoting the X and Y positions of residues contributing to PAS-C folds. **G.** Structural alignment of the PAS-C domain output from RosettaFold and ESMFold with bacterial FixL. **H.** Structural alignment of PAS-C domain output from RosettaFold and ESMFold with the human Arnt PDB structure.

Intrigued by the location of residues that contribute to the structural elements of the PAC motif, Gβ, Hβ, and Iβ sheets, we investigated whether these residues align well with SMART predicted PAC motifs from spotted gar, West African lungfish, passerine birds, and Himalayan leaf-nosed bats (Figure S4). While the SMART algorithm did not predict the presence of PAC-motifs for humans or rifleman PASK, aligning them with species where the PAC motif was predicted showed high sequence conservation. Furthermore, structural alignment using the hPASK-AF2 structure as a template confirmed both sequence and structural similarities, indicating that these sequences in human and rifleman PASK are unannotated PAC motifs that co-assemble into the PAS-C domain, despite being separated by a variable linker length (Figure 3D).

To further scrutinize the PAS-C domain assembly models, we assessed the Predicted Aligned Error (PAE) matrix to analyze the interaction between the N-terminal PAS fold (residues 345-402) and C-terminal PAC motif (residues 888-931) in human PASK. As expected, for rifleman PAS-C, the PAE scores between the N-terminal PAS and C-terminal PAC residues were low because the PAS-C domain was formed by a contiguous sequence (Figure 3E). Notably, in human PASK, the PAE scores for the PAS-C domain were also low (average PAE score of 4 Å), similar to the rifleman PASK PAS-C domain (Figure 3F). This indicated well-modeled long-range interdomain interactions driving the assembly from two regions separated by nearly 500 amino acids.

To identify a structural ortholog of the PAS-C domain, we employed the independent model validation workflow described in Figure 2F. We combined amino acids contributing to the N-terminal PAS-C (aa345-402) with the C-terminal PAC motif (aa888-931) of human PASK and queried this combined sequence against the CATH structural classification database. The CATH database identified the PAS-C motif as belonging to PAS family structure cluster 3.30.450.20/2, which is closely related to the sensor protein from *Bradyrhizobium japonicum* FixL functional family, with an E-value of 9.8e-4.

Similarly, analysis of independent PAS-C domain structures obtained from either RosettaFold or ESMFolds using FoldSeek identified FixL from *Bradyrhizobium japonicum* (Figure 3G) and the PAS-B domain from ARNT as the two closest structural matches (Figure 3H). Based on these comprehensive analyses, we conclude that the PAS-C domain in most metazoan PASK is derived from bacterial oxygen sensors and might be assembled through unique intramolecular interactions between PAS and PAC fragments separated by a variable-length linker.

### The improved HMMER profile broadened the identification of the PAC motif-linked PAS domain across the metazoa

Our analysis revealed that the HMMER-based SMART algorithm inconsistently identified PAC motifs in PASK proteins from various metazoan species. This inconsistency likely stems from the algorithm’s reliance on a specific sequence identity threshold set by HMMER. Variations in the PAC domain sequence, often due to genetic drift, can lead to unpredictable PAC motif predictions in metazoan PASK, as certain residues fluctuate around the prediction threshold. This is supported by evidence that, when searching for PAS domain-containing proteins using the HMMER profile for PAS domains (SMART accession number: SM00091), SMART identified approximately 100 PAS domain-containing proteins in the human proteome. In contrast, a SMART-based search using the PAC-motif profile (accession number: SM00086) returned only 79 PAS domain-containing proteins and failed to identify PASK in the human proteome as a PAS domain-containing protein. This suggests that the PAC motif profile used by SMART/HMMER is not sufficiently comprehensive to accurately identify all PAS domain-containing proteins.

To improve PAC domain identification, we generated an enhanced HMMER3 (termed i-HMMER3, Supplementary Dataset 1) profile by aligning known PAC domain sequences from prokaryotes and eukaryotes with PAC domain sequences from 18 species of PASK, where SMART identified the PAC domain (Figure S4, Figure S6A-B). We compared the performance of the PAC profile from i-HMMER3 with that of SMART and Prosite (Prosite matrix, accession number: PRU00141) to identify PAS domain-containing proteins in the human proteome. As indicated earlier, the SMART-based PAC profile identified 79 PAS domain-containing proteins (including isoforms and splice variants) (Figure S6C) linked to the PAC domain, whereas Prosite identified only 26, including false negatives (Figure S6C). i-HMMER3 successfully identified all PAS domain-containing proteins detected by SMART and uncovered 20 additional splice variants and tissue-specific isoforms of bHLH transcription factors (e.g., O00327-8 or Bmal1-8) in the human database (Figure S6C), with PAS and PAC domains identified in tandem. When scanned against the metazoan proteome, i-HMMER3 identified nearly 15,000 PAS domain-containing proteins compared to SMART’s 9,000 proteins identified by SMART (Figure S6D).

Additionally, i-HMMER3 successfully identified human PASK and its splice variants as PAS domain-containing proteins. i-HMMER3 identified the PAC domain in all metazoan PASK proteins, located consistently 63-69 amino acids upstream of the kinase domain and 16-650 amino acids on the C-terminal side of the truncated PAS-C domain (Figure S7A). These findings confirmed the identification and location of the PAC domain in the metazoan PASK and expanded the automatic annotation of PAS-PAC tandem domains across metazoans.

Our analysis showed that in most metazoan PASK, the N-terminal PAS-fold and C-terminal PAC-folds, which form the PAS-C domain, are separated by a linker of variable length (486 amino acids in humans; see Figure S7B). According to our model, these distinct folds together form a functional PAS-C domain through long-range intramolecular interactions (Figure S7C).

### Interdomain interactions and cross-linking studies validated the PAS-C Domain assembly model

To experimentally validate the AlphaFold structure of the PAS-C domain and its unique mode of assembly, we generated a structure-based sequence alignment of the human PAS-C domain with structurally related PAS domains from the CATH database to identify functionally conserved residues (Figure S8). The metazoan PAS-C domain of PASK contains conserved residues at key positions in the FixL and other PAS family members. For instance, the human PAS-C domain residues I357 in the Bβ strand, E375 in the Dα helix, and Y391 are conserved in the metazoan PASK and other PAS domain-containing proteins. In most terrestrial PASK, the heme-coordinating histidine of bacterial FixL in the Fα helix is replaced by aspartate (D) or glutamate (E) (Figure S8, highlighted in the red box). Conversely, most actinopterygians, including spotted gars and milkfish, either retain histidine or substitute it with other amino acids capable of coordinating heme such as cysteine or serine (Figure S8). However, other heme-interacting residues in FixL (R206 and H214) are not conserved in fish, which may indicate a gradual loss of heme-binding capabilities of the PAS-C domain of PASK. Consistent with this, we observed the emergence of Asp (D) in lungfish, the closest living ancestor of tetrapods linked to the water-to-land transition (Figure S8).

To identify the critical residues in the PAC domain that engage in PAS domain assembly, we specifically aligned the PAC domain of mammalian PASK with PAC domain sequences in the existing SMART/HMMER/PDB databases (Figure S9A-B). We identified the PAC domain of the plant histidine kinase Phytochrome C (PhyC) as a close structural homolog of the mammalian PAC domain of PASK. When comparing the structural assembly and interaction of the C-terminal beta-strands (Gβ, Hβ, and Iβ) which form the PAC domain in the metazoan PASK (Figure S8 and S9) with the N-terminal PAS fold between human PASK, bacterial FixL, and plant PhyC, we observed a strongly conserved H/R-R-D-G sequence in PASK (900H-R-D-G903), FixL (226R-R-D-G229), and PhyC (830H-R-D-G833) in the loop region between the Gβ and Hβ sheets (Figure S8 and S9). In the AlphaFold structure of PASK, R901 within the human PAC fold forms a salt bridge with the highly conserved E375 residue within the Dα helix of the PAS fold (Figure 4A). The Arg-Glu salt bridge is also observed in the crystal structure of FixL (R227-E182, PDB ID:1DP6)[23] (Figure 4A), the AlphaFold model of the Phytochrome C PAS-B domain (R831-E778), and the experimentally derived structure of the PAS-B domain of ARNT (PDB: 2b02), where K432 in the loop between the Gβ and Hβ sheets forms a salt bridge with E390 of the Dα helix (Figure 4A). Thus, a sequence-structure comparison of the PASK PAC motif with other PAS domain-containing residues identified residue pairs conserved across PAS domain structures from prokaryotes to humans.

**Figure 4.**
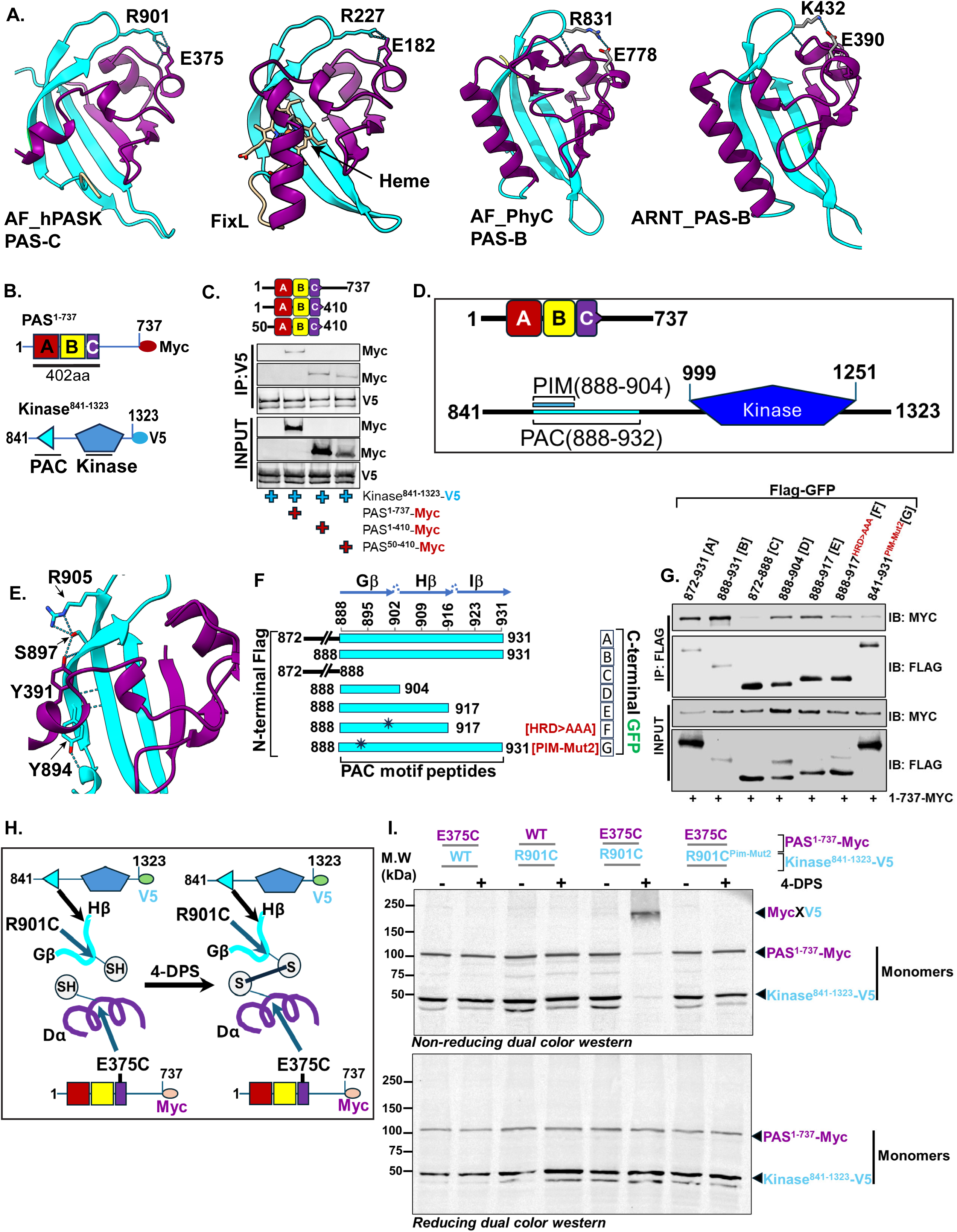
Experimental Validation of the PAS-C Domain Assembly in Human PASK. **A.** Comparison of the human PAS-C structural model with the FixL PAS domain (PDB ID: 1DP6), the AlphaFold model of the *Arabidopsis thaliana* PAS-B domain, and the human Arnt-PAS-B domain (PDB: 2b02). **B.** Domain architecture of two fragments of PASK used to assess the interaction between two regions of PASK. Top: PAS-A, PAS-B, and PAS-C domains comprise 400 amino acids within the first 1-737 fragment. Bottom: Domain layout in the C-terminal fragment, 841-1323, showing the location of the PAC motif and the serine/threonine kinase domain. **C.** Co-expression of the indicated N-terminal and C-terminal fragments in HEK293T cells, with their interaction measured by co-immunoprecipitation followed by western blotting using the indicated antibodies. **D.** Domain schematic illustrating the overlap between sequences previously described as the PAS-Interacting Motif (PIM) and PAC motif in the C-terminal region of PASK. **E.** AlphaFold model of the PAS-C domain of PASK showing the interaction between S894-Y391-R905 of the PAS-C domain. **F.** Mapping of the minimal PAC motif requirement for interaction with the PAS^1-737^ fragment. The peptide PAC motif sequence shows the location of individual β-sheets of the PAC domain, and various Flag-GFP-tagged peptides were used to determine the interaction of the PAC motif with the N-terminal fragment. A star in the bottom two peptides, 888-917 and 888-931 peptides, indicates an approximate location of mutations used to probe their functional roles. **G.** Co-immunoprecipitation of Flag-GFP tagged WT or mutated PAC motif peptides with Myc-tagged-PAS^1-737^ fragment from HEK293T cells. Peptides were purified from cells using an anti-FLAG antibody, and the co-precipitation of Myc-tagged PAS^1-737^ fragments was determined by western blotting using the indicated antibodies. **H.** Schematic of the VivoX experimental setup. **I.** VivoX analysis of R901C and E375C interactions. Cells expressing the indicated plasmids were treated with 180µM 4-DPS (+) or DMSO (−). Cell extracts were prepared and separated by SDS-PAGE under reducing or non-reducing conditions. The proteins were detected using the indicated antibodies.

We recently demonstrated that the intramolecular interaction in PASK involves the C-terminal region (aa 841-1323, termed Kinase^841-1323^) and N-terminal region (aa ^1-737^, termed PAS^1-737^) of PASK (Figure 4B)[24]. This interaction is mediated by a short, linear interacting motif, which we termed the PAS-interacting motif (PIM) (Figure 4C-D)[24]. An improved understanding of the PAS^1-737^ region through this study suggests the presence of two PAS domains (PAS-A and PAS-B) and a shorter truncated PAS-C domain within the first 402 amino acids (Figure 4B). Thus, we tested whether the Kinase^841-1323^ fragment can associate with a minimum region containing PAS-A, PAS-B, and truncated PAS-C domains in cells. Consistent with our previous results, Kinase^841-1323^ could immunoprecipitate the PAS^1-737^ fragment of PASK (Figure 4C). Furthermore, the Kinase^841-1323^ fragment also interacted with PAS1-410 (lacking residues 41^1-737^) and PAS50-410 (lacking the first 50 amino acids and residues 41^1-737^), indicating that the intramolecular interaction is primarily targeted to one of the three PAS domains (Figure 4C). Our previous results suggested that residues 888-915 within the Kinase^841-1323^ fragment are required for intramolecular interaction with PAS^1-737^[24]. Within this region, we identified a stretch of residues E888-L904, which we termed the PAS interacting motif (PIM), as necessary and sufficient for interaction with the PAS^1-737^ fragment [24]. Furthermore, residues Y894 and S897 within the PIM motif are crucial for interaction with the N-terminal region of PAS^1-737^[24]. Interestingly, the PIM sequence overlapped with the PAC motif of human PASK (E888-D931) (Figure 4D). In the AlphaFold structure of human PASK, S897 forms hydrogen bonds with the highly conserved Y391 in the Fα helix of the PAS fold and R905 in the Gβ of the PAC domain, thus anchoring the Fα helix to the β-sheets of the PAC fold (Figure 4E). To precisely map the PAC domain boundary that associates with the N-terminal PAS domains, we generated a series of overlapping Flag-GFP fused peptides (Figure 4F) that encompass Gβ (aa 888-904), Gβ and Hβ (888-917), and Gβ, Hβ, and Iβ (888-931) sheets. We co-expressed these fusion proteins with PAS^1-737^ and measured their protein-protein interactions by co-immunoprecipitation. As shown in Figure 4G, the entire PAC domain encompassing residues 888-931 exhibited the strongest interaction with the PAS^1-737^ fragment, while region 888-904 was necessary and sufficient to interact with PAS^1-737^, in agreement with our recent results [24]. Mutations in Y894 and S897 to Ala (PIM-Mut2) [24] markedly reduced the interaction between the PAC-fold and PAS^1-737^ regions. Furthermore, mutations in the conserved H/R-R-D-G loop between Gβ-Hβ (^900^H-R-D-G^903^, Figure 4A) and HRD>AAA disrupted the interaction between the PAC motif and PAS^1-737^ (Figure 4G), suggesting that these residues are also important for the intramolecular association between the PAC fold and PAS^1-737^.

To directly test the hypothesis that the PAS-C domain is reconstituted in cells through intramolecular assembly of the N-terminal PAS fold (aa 345-402) and PAC fold (aa 888-931) domains, we utilized the VivoX method to probe the juxtaposition and interaction between R901 and E375 (Figure 4A) [25]. Guided by the AlphaFold model of the PAS-C domain, we mutated R901 and E375 to cysteine residues in the C-terminal- and N-terminal fragments, respectively (Figure 4H). When co-expressed in cells, R901C and E375C would form disulfide cross-links upon addition of 4-DPS if they are juxtaposed, resulting in migration of the adducts under non-reducing SDS-PAGE conditions (23). Excitingly, we noticed strong disulfide adduct (MycXV5) formation between Kinase^841-1323(R901C)-V5^ and PAS^1-737-(E375C)-Myc^ under non-reducing conditions in 4-DPS-treated cells, which were resolved under reducing conditions. Interestingly, disruption of the PAS-C domain assembly due to Y894 and S897 mutations to alanine (PIM-Mut2) completely disrupted disulfide adduct formation between R901C and E375C (Figure 4I), indicating a loss of juxtaposition between these residues. These results and our structural analyses confirm and validate the AlphaFold model of PAS-C domain assembly and suggest that the PAS-C domain of PASK is assembled through intramolecular interactions of disconnected PAS and PAC motifs.

### The Intramolecular assembly of PAS-C facilitates catalytic core stabilization by remodeling the quaternary structure of PASK

The unique assembly of the PAS-C domain led us to investigate its role in human PASK’s catalytic function. Rifleman PASK, being shorter and featuring a contiguous PAS-C domain, presented an ideal model to explore the functional implications.

First, we assessed whether rifleman PASK could functionally replace mammalian PASK. Both human and mouse PASK are essential for muscle cell differentiation and regeneration. Silencing mouse PASK in cultured muscle cells inhibited this process (Figure 5A-B). Remarkably, introducing rifleman PASK rescued this defect, confirming functional conservation across species (Figure 5A-B).

**Figure 5.**
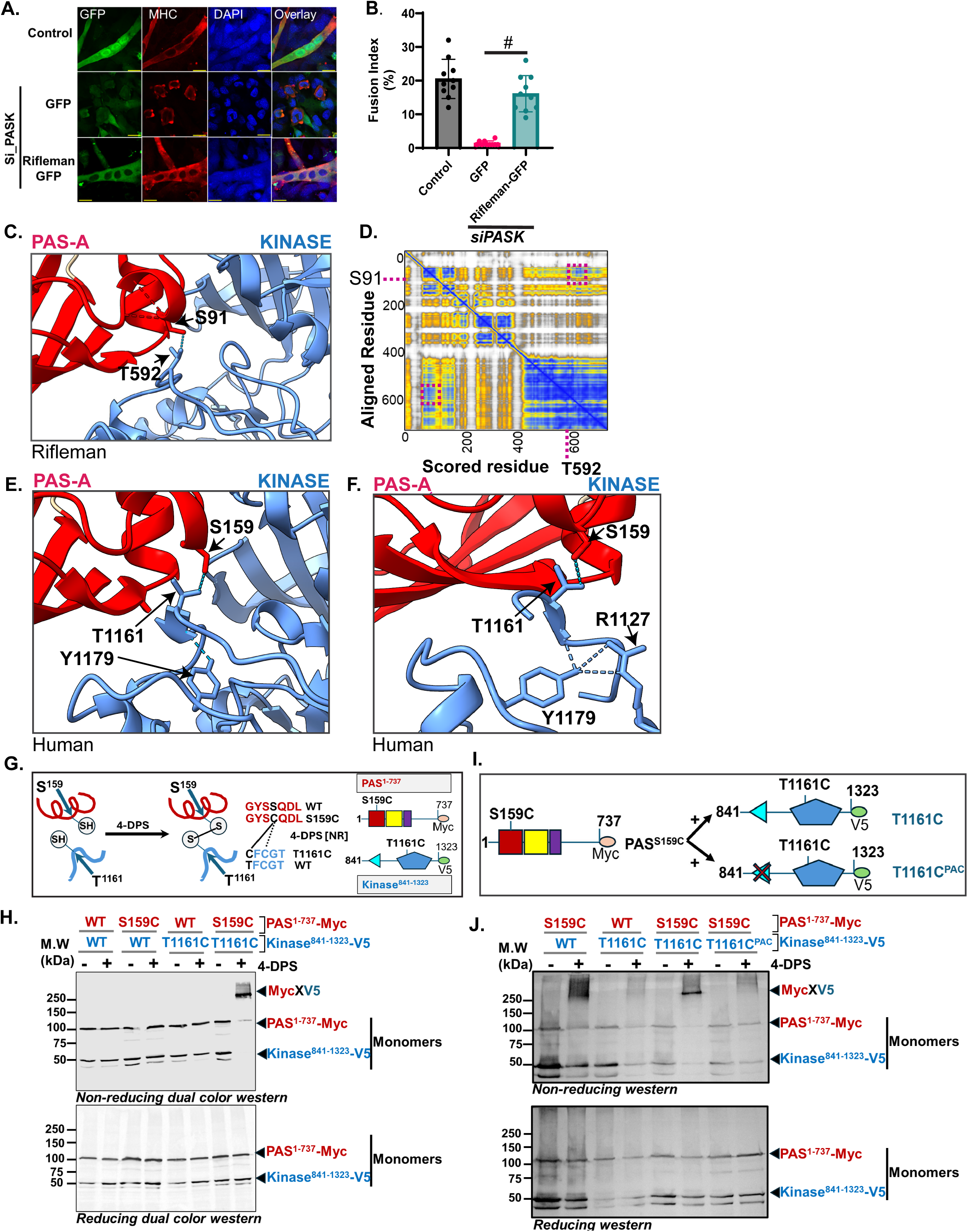
Intramolecular Assembly of the PAS-C Domain Stabilizes the Catalytic Core of PASK. **A.** Mouse PASK was silenced using smart-pool siRNA duplexes in proliferating C2C12 myoblasts. 24 hours after transfection, the C2C12 cells were infected with retroviruses carrying either GFP or GFP-Rifleman PASK. 48 hours after infection, myogenesis was induced by adding 2% horse serum and differentiation was allowed to proceed for an additional 48 h. The cells were fixed and myogenesis was detected using an anti-myosin heavy chain antibody (MF20). Scale bars = 40µM. **B.** Quantification of the fusion index indicating myogenesis efficiency from the experiment in **A.** Each circle corresponds to fusion index from biological replicates. N=10. Error bars = ± S.D. #*P<0.001.* **C.** Interdomain interactions between PAS-A (red) and the kinase (blue) domains of rifleman PASK. S91 of PAS-A is engaged in a hydrogen-bonding interaction with T592 of the activation loop motif of the PASK kinase domain. **D.** Position Error Matrix (PAE) for the AlphaFold structure of rifleman PASK showing the positional score between S91 of the PAS-A domain and T592 of the kinase domain (magenta rectangle). **E.** Evidence of interaction between S159 of human PAS-A and T1161 of the human Kinase domain in the AlphaFold structure. **F.** Catalytic loop stabilization by R1127-Y1179-T1161-S159 interactions in AlphaFold structure of human PASK. **G.** Schematic of the VivoX experimental setup to test the interaction between S159 and T1161. The αD helix of the PAS-A domain containing S159 is depicted in red, and the activation loop of the PASK kinase domain is in blue. WT and cysteine-mutated sequences for each residue are shown. Note the presence of a naturally occurring cysteine residue in the PASK activation loop sequence in the WT PASK sequence (Cys1163). The panel to the right shows the constructs used in the VivoX experiment to test the interaction between the PAS-A domain (red) and kinase domain (blue). **H.** Inter-residue cross-linking (VivoX) analyses of PAS-A and Kinase domain interactions. HEK293T cells expressing Myc-tagged WT or S159C mutated PAS^1-737^ along with WT or T1161C mutated V5 tagged 841-1323 constructs were treated with 180µM 4-DPS (+) or DMSO (-) for 20 min before cell lysis and analyzed by non-reducing (TOP) and reducing (Bottom) western blot analysis, followed by a combination of Myc and V5-tagged antibodies on the same membrane. Notice a faint smearing in lane 2, perhaps because of long-range cross-linking between S159C and Cys1163. **I.** Schematic of the experimental setup to test the contribution of the human PAC domain in bringing PAS-A and Kinase domain into proximity. **J.** Interresidue cross-linking (VivoX) analyses of PAS-A and Kinase domain interactions with or without mutations in PA**C.** HEK293T cells expressing combinations of WT or S159C mutated PAS^1-737^ with WT or PAC mutated 841-1323T1161C constructs were treated with 180µM 4-DPS (+) or DMSO (-) for 20 min before cell lysis, and analyzed by non-reducing (TOP) and reducing (Bottom) western blot analysis, followed by combined Myc and V5-tagged antibodies on the same membrane.

This conservation is notable given the evolutionary distance, differing protein lengths, and variations in PAS-C formation (Figure 3A-B). Such functional conservation suggests a shared mechanism of action, likely involving a common quaternary structure arrangement. To explore this, we examined the quaternary structure of rifleman PASK by computing Predicted Aligned Error (PAE) scores. While the rifleman kinase domain is not generally juxtaposed with any of the PAS domains, based on elevated PAE scores, we observed a significantly lower PAE score between the residue pair S91 in the Dα helix of the PAS-A domain and T592 in the kinase domain of rifleman PASK (PAE score = 5.9) (Figure 5C-D, purple box), indicative of close juxtaposition between specific regions of PAS-A and kinase domains of rifleman PASK. Consistent with that, in rifleman PASK AlphaFold structure, S91 of PAS-A domain hydrogen bonds with T592 of the Kinase domain.

T592 of rifleman PASK and the corresponding T1161 of human PASK are in the activation loop of the catalytic domain. In most serine/threonine protein kinases, the first threonine of the activation loop sequence (typically T-F/Y-C-G-T) is phosphorylated by upstream kinases to induce the necessary conformational changes for substrate peptide binding and catalysis [26]. We previously demonstrated that phosphorylation of threonine within the activation loop (T1161) is not required for the catalytic activity of human PASK and that the open catalytic conformation is achieved via side-chain-mediated interactions [13]. For instance, the in unphosphorylated state, activation loop T1161 of human PASK interacts with Y1179 via main-chain atoms (Figure 5E). Through this interaction, Y1179 helps to stabilize the catalytically active conformation through triangulation with the highly conserved R1127 (within the HRD motif found in most kinase domains) and T1161 (Figure 5F). However, it remains unclear how the side chain of T1161 is coordinated to stabilize the activation loop. In most activation-loop phosphorylation-dependent protein kinase structures, the activation-loop serine or threonine residue is stabilized by the interaction with Arg from the αC-helix of the kinase domain upon phosphorylation [27]. As T1161 of PASK is not phosphorylated in cells, and the αC-helix of PASK is unusually truncated [13], the mechanism stabilizing the T1161 side chain remains unclear. Interestingly, AF2 predicted a highly similar quaternary structure arrangement between human and rifleman PASK, where S159 of human PAS-A formed a hydrogen bonding interaction with the activation loop T1161 of the Kinase domain (Figure 5E-F). Thus, we wondered whether S159 of the PAS-A domain could stabilize T1161 of the activation loop via interdomain association between PAS-A and the kinase domain (Figure 5E-F).

To test this hypothesis, we employed the VivoX cross-linking assay described earlier (Figure 4H-I). We generated S159C and T1161C mutations in the N-terminal (PAS^1-737^) and C-terminal (Kinase^841-1323^) fragments, respectively, and expressed them in HEK293T cells in various combinations of mutant and WT versions of the fragments. When PAS^1-737(S159C)^ was expressed with the Kinase^841-1323(WT)^ fragment, we noticed slight but inconsistent smearing of the gel shift under non-reducing conditions, perhaps due to the weak, opportunistic disulfide bonds between S159C and Cys1163 within the activation loop (Figure 5G, dotted line, and Figure 5J). The expression of the Kinase^841-1323(T1161C)^ fragment with PAS^1-737(WT)^ did not show any noticeable gel mobility shift under non-reducing conditions. Remarkably, when PAS^1-737(S159C)^ was co-expressed with Kinase^841-1323(T1161C)^, we noticed strong MycXV5 adduct formation under non-reducing conditions, which was resolved under reducing gel conditions (Figure 5H). These results confirm that S159 in the PAS-A domain is in structural proximity and is likely an interacting partner of T1161 in the kinase domain in cells.

Intrigued by the consistent amino acid spacing from the end of the PAC motif and the beginning of the kinase domain (Figure S7A-B) and the intramolecular assembly of PAS:PAC folds that generate the PAS-C domain (Figure 4), we wondered whether PAS-C domain assembly might tug on the kinase domain to bring it into proximity to the PAS-A domain, thereby organizing the overall quaternary structure (Figure S7C). To test this, we generated a PAS-C assembly defective PAC mutant (Y894AS897AR901A) in the T1161C-mutated Kinase^841-1323^ fragment (Kinase^841-1323 (T1161C)/PACmut^) (Figure 5I). As expected, co-expression of PAS^1-737(S159C)^ with Kinase^841-1323(T1161C)^ resulted in strong MycXV5 adduct formation under non-reducing conditions. However, disrupting PAS-C domain assembly via PAC mutations abolished the MycXV5 adducts between PAS^1-737(S159C)^ and Kinase^841-1323(T1161C)/PACmut^ (Figure 5I-J). These results suggest that intramolecular assembly of the PAS-C domain organizes the quaternary structure of PASK to facilitate its interaction with the kinase domain, stabilizing the activation loop Thr1161, and providing an avenue for its regulation.

### The Signal-Regulatable quaternary structure remodeling of PASK via PAS-C Domain assembly

Previous structural studies have proposed that the metabolite-binding PAS-A domain could influence the catalytic functions of PASK via direct interaction with its kinase domain [14]. We experimentally confirmed the direct pairing between the PAS-A and Kinase domain, which is facilitated by the PAS-C domain assembly (Figure 5I-J). However, we wondered whether regulated PAS-C domain assembly could modulate PAS-A and kinase domain juxtaposition, potentially leading to dynamic remodeling of the catalytic loop T1161.

To test this hypothesis, we assessed whether PAS-C assembly is signal-regulated in the cells. Due to limitations related to the unequal expression levels of separately overexpressed N- and C-terminal fragments in cell-based assays for protein-protein interaction measurements, we engineered a full-length PASK protein incorporating an internal TEV protease cleavage site between its N-terminal and C-terminal regions, separating PAS and PAC folds of the PAS-C domain, as depicted in Figure 6A. Using this construct, we tested whether the N-terminal and C-terminal regions of PASK maintained their association following TEV protease-mediated cleavage of the loop region connecting the two domains, indicative of stable intramolecular domain formation. As indicated in Figure 6B, the association between the N- and C-terminal fragments persisted after TEV cleavage (lane 2) in the WT-Flag-TEV-Kinase-V5 proteins, reinforcing evidence for the intramolecular interaction between these regions. Conversely, mutation of the PAC motif residues resulted in the loss of association between the N- and C-terminal fragments, supporting the role of the PAC fold in facilitating intramolecular assembly.

**Figure 6.**
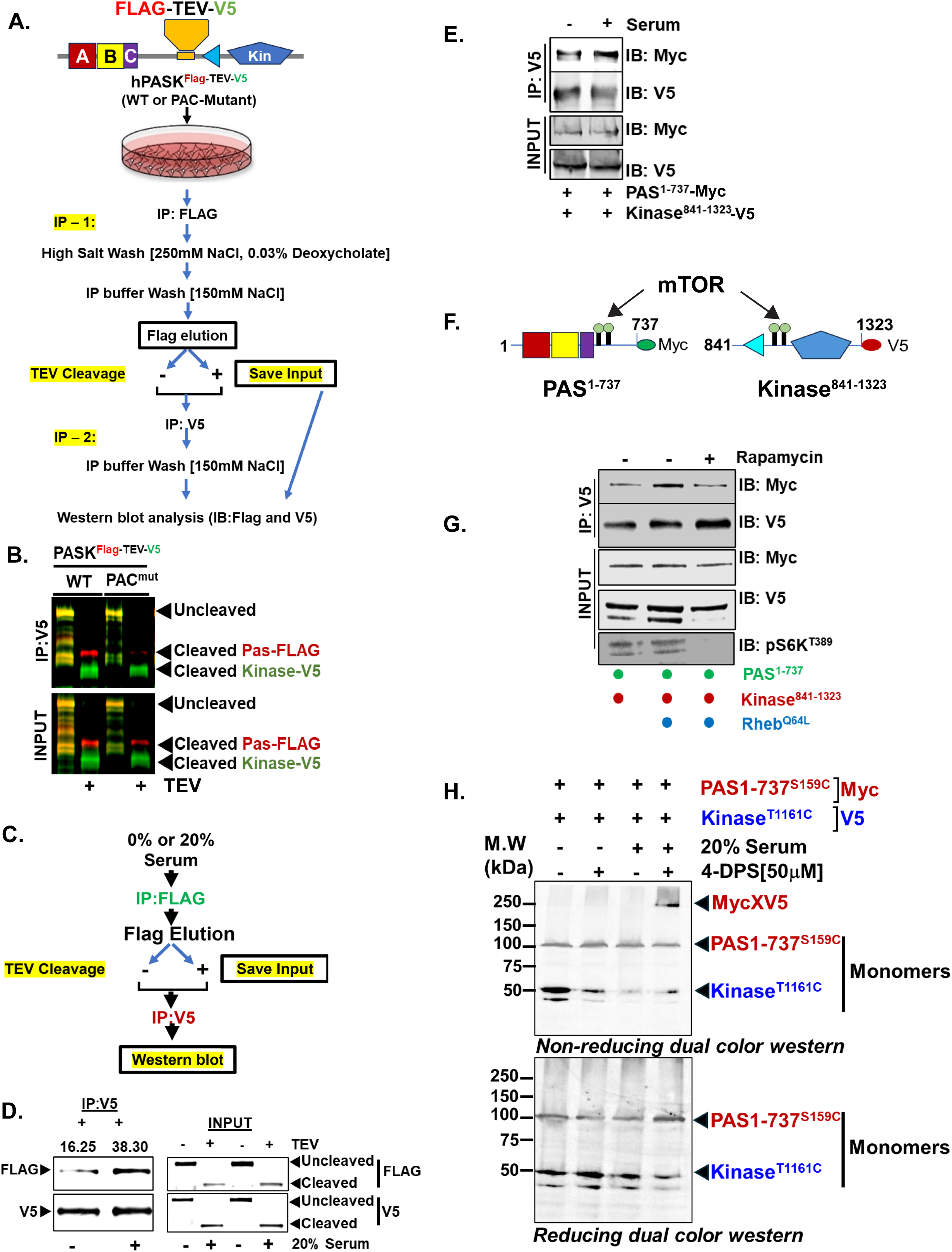
Serum Stimulation Organizes Quaternary Structure of PASK by Promoting Intramolecular Assembly. **A.** Assessment of intramolecular interaction in PASK using an internal TEV protease system. **B.** HEK293T cells expressing WT or PAC-mutated hPAS-Flag-TEV-V5-Kinase were lysed and processed according to the experimental layout in **A.** V5-tagged proteins were purified before and after TEV cleavage, and the extent of co-associated Flag-PAS (1-735) was determined by western blotting. **C.** Schematic to test the effect of serum stimulation on intramolecular interactions between N-terminal PAS domains and C-terminal PAC-kinase domains using a TEV protease cleavage system. **D.** HEK293T cells expressing hPAS-Flag-TEV-V5-Kinase were serum-starved overnight, followed by serum stimulation for 20 mins with 20% fetal bovine serum (FBS). Cells were lysed and processed as shown in Figure **A.** V5-tagged proteins were purified and the extent of co-associated Flag-PAS was detected by western blotting. **E.** HEK293T cells expressing Myc-PAS^1-737^ and V5-Kinase^841-1323^ were serum-starved overnight, followed by serum stimulation with 20% serum for 20 mins. Cells were lysed and proteins were purified using anti-V5 beads. Co-immunoprecipitation of FLAG was detected by western blotting. **F.** Domain schematic showing the location of mTOR phosphorylation sites in PAS^1-737^ and Kinase^841-1323^. **G.** HEK293T cells expressing Myc-PAS^1-737^ and V5-Kinase^841-1323^ were serum-starved overnight, followed by serum stimulation with 20% serum for 20mins in the presence or absence of 100nM Rapamycin. Cells were lysed, and proteins were purified using anti-V5 beads. Co-immunoprecipitation of Flag was detected via western blotting. **H.** Serum regulation of interdomain interactions was performed using the VivoX method. **D.** HEK293T cells expressing PAS^1-737(S159C)^ and Kinase^841-1323(T1161C)^ were serum-starved overnight. The next day, the cells were treated with 50µM 4-DPS (+) or vehicle (DMSO, -) with or without 20% FBS for 20 min. Cells were lysed, protein extracts were prepared under reducing or non-reducing conditions, and SDS-PAGE was performed to detect Myc-V5 adduct formation.

Interestingly, in an analogous experiment, when acutely stimulated with serum, we observed a 2-fold increase in the association between the N-terminal PAS^1-737^ and the C-terminal PAC-kinase domain-containing fragment (Figure 6C-D). We also confirmed the serum-stimulated interaction between the N-terminal and C-terminal regions in a co-expression system (Figure 6E) using a Myc-tagged PAS^1-737^ fragment (N-terminal) and a V5-tagged Kinase^841-1323^ fragment (C-terminal). Since the interactions between the N- and C-terminal fragments are mediated through PAC motif sequences (Figures 4G-I and 6B), it is possible that the signal-regulated assembly of the PAS-C domain enhances serum-stimulated interactions between the N-terminal (PAS^1-737^) and C-terminal (Kinase^841-1323^) fragments.

We previously reported the catalytic activation of PASK via the mTOR complex 1 (mTORC1)-mediated phosphorylation of residues S640 and T642 within the PAS^1-737^ region and S949, T953, and S957 within the C-terminal Kinase^841-1323^ region of PASK juxtaposed with the PAC motif (Figure 6F, green circles) [11]. Since these phosphorylation motifs are outside of well-defined domain boundaries, it remains unclear how mTORC1-mediated phosphorylation could stimulate PASK catalytic activity. Given the role of the PAC motif in PAS-C domain assembly (Figure 4) and stabilization of the PASK catalytic core via S159-T1161 interactions (Figure 5J), we speculated that mTORC1 phosphorylation enhances the interactions between the N-terminal and C-terminal regions of PASK and catalytic activation. In agreement with this notion, acute serum-stimulation resulted in a robust interaction between the PAS^1-737^ and Kinase^841-1323^ regions of PASK, which were diminished by pretreatment with the mTOR inhibitor rapamycin (Figure 6G).

As mTORC1 stimulation led to increased PASK catalytic activity, we wondered if serum stimulation remodels the quaternary structure of PASK through PAS-C domain assembly, thereby potentially linking mTORC1 signaling with catalytic core stabilization (Figure 5H-J). Consequently, we investigated whether serum-stimulated assembly of the PAS-C domain results in a stronger interaction between PAS-A and the activation loop of the kinase domain. To test this hypothesis, we repeated the VivoX experiment using PAS^1-737^ (S159C) and Kinase^841-1323(T1161C)^ fragments in the presence or absence of acute serum stimulation in mammalian cells. To enhance the sensitivity and specificity of the VivoX reaction, we reduced the concentration of 4-DPS to 50 µM for this experiment. We co-expressed Myc-tagged PAS^1-737 (S159C)^ and V5-tagged Kinase^841-1323 (T1161C)^ fragments in HEK293T cells and analyzed serum-stimulated adduct formation at a low concentration of 4-DPS. Surprisingly, serum starvation led to a near-complete loss of MycXV5 adduct formation between PAS^1-737(S159C)^ and Kinase^841-1323(T1161C)^, phenocopying the effect of PACmut on S159C-T1161C interaction (Figure 5I-J). Thus, serum withdrawal is likely leading to the disassembly of the quaternary structure arrangement critical for the PAS-A:Kinase domain interaction (Figure 6H). In contrast, acute serum stimulation robustly induced MycXV5 adduct formation, even at low 4-DPS concentrations, between S159C of PAS-A and the activation loop T1161 of the kinase domain (Figure 6H), indicating increased PAS-A:Kinase domain tethering to enhance the stability of the catalytic core in response to serum. Combined with the serum-stimulated increase in the interaction between PAS^1-737^ and Kinase^841-1323^ (Figure 6D-G), which is dependent on the PAC sequence (Figure 6A) and PAS-C domain assembly (Figure 4I), our results suggest a unified model of PASK activation by serum through PAS-C domain reconstitution, which remodels the quaternary structure of PASK for full catalytic activation. These results indicate that signaling inputs, such as mTORC1-mediated phosphorylation within the linker region, dynamically regulate PAS-C domain assembly, leading to the catalytic stabilization of the PASK activation loop via quaternary structure rearrangement.

## Conclusion

Through the phylogeny-guided domain architecture analysis and structure comparison approach, we identify a novel mechanism of PAS domain assembly in metazoan PASK. We show that nutrient signaling influences activation loop stabilization in the metazoan PASK through quaternary structure rearrangement by stabilizing the PAS-C domain assembly. As an important protein kinase that regulates stem cell differentiation and metabolic decisions, the precise control of its catalytic activity is warranted. Our studies demonstrated that PASK has evolved a unique mode of domain assembly to achieve this (Figure 7A-B). Broadly, our approach of comparative structural phylogenetic analysis could provide a framework for extracting physiologically relevant quaternary structural information from deep-learning-based structural models.

**Figure 7.**
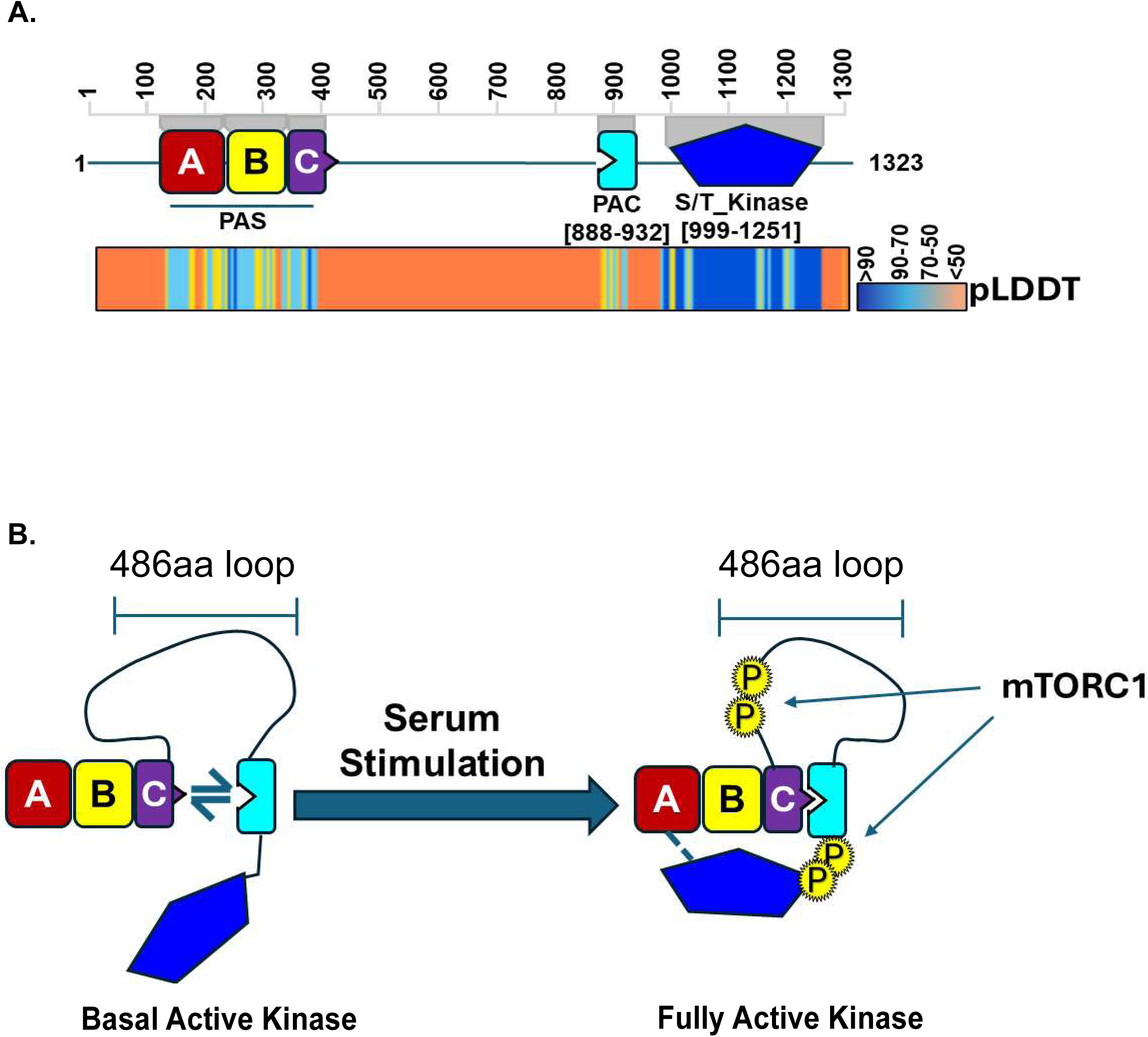
An integrated model of the PAS-C domain assembly and catalytic activation of PASK through quaternary structure remodeling. **A.** Structure and i-HMMER-based revised domain architecture of PASK overlaid with pLDDT scores from the human AlphaFold structure, matching structural elements. **B.** Our model of PASK activation in response to serum stimulation following mTORC1-mediated phosphorylation. Under steady-state conditions, the PAC domain is partially unfolded from the globular PAS secondary structure of the PAS-C domain (see the discussion for other examples). Under these conditions, PASK is basally active. Serum stimulation stabilizes PAS-C assembly, increasing the juxtapositioning of the kinase domain with the PAS-A domain, leading to coordination of activation loop T1161 side chain with S159 of the PAS-A domain.

## Discussion

In this evolutionary structural biology study, we revealed a novel, regulatable principle of protein folding that relies on the long-range intramolecular assembly of two PAS protein fold subdomains in the nutrient-sensing kinase PASK: the N-terminal PAS fold and the C-terminal PAC fold. We demonstrated that serum stimulation remodels the quaternary structure of PASK via the intramolecular assembly of a third PAS domain, PAS-C, thereby linking protein folding with the catalytic activation of PASK. Building on these findings, we propose that PAS domains exhibit greater dynamism in their tertiary structure assembly, and that the loop sequences bridging PAS folds and PAC folds could be key targets of signaling pathways.

### Phylogeny-guided domain architecture and structure comparison workflow

We developed an analytical workflow that combined phylogenetic analysis with domain mapping and quaternary structure analysis on an evolutionary scale to elucidate the presence of three PAS domains in PASK. The first two PAS domains of PASK, PAS-A, and PAS-B span approximately 100 amino acids each and show a canonical 5α/5β secondary structure arising from a contiguous amino acid sequence that folds PAS and PAC together (Figure 2H). The third PAS domain, PAS-C, also contains a 5α/5β structure; however, its tertiary structure is uniquely assembled through intramolecular interactions between the PAS and PAC folds (Figure 3). This conclusion is supported by our extensive analysis employing computational protein modeling, sequence-structure comparisons, large-scale structure-based sequence alignments across bacterial and eukaryotic PAS domain-containing proteins, and detailed experimental analysis of the structural model at the resolution of individual residues. Our phylogeny-guided domain architecture analysis identified a single PAC domain in approximately 18 species of the metazoan PASK. Through co-immunoprecipitation and in vivo cross-linking assays (VivoX), we were able to show interactions between residues in PAS and PAC folds, leading to a novel understanding of PAS domain folding through long-range intramolecular interactions. This analysis confirmed the success of our structural phylogeny approach in validating the quaternary structure of deep learning-based protein structure models.

### PAS-C Domain Assembly and Its Role in Catalytic Core Stabilization

In every metazoan species studied, the PAC domain of PASK is positioned 63-69 amino acids upstream of the kinase domain, a pattern not observed for the loop that connects the PAS and PAC folds (Figure S7A-B). This observation led us to hypothesize that assembly of the PAS-C domain influences the quaternary structure of PASK. Although AlphaFold models are generally reliable for predicting local tertiary structures when pLDDT scores exceed 70, the biological relevance of the predicted quaternary structures from AlphaFold remains uncertain. Our analysis suggests that comparison of AlphaFold-derived quaternary structures of both related and distant species can increase confidence in these predictions. For instance, we found remarkable similarities in the quaternary structures of rifleman and human PASK, despite significant differences in the PAS-C assembly mode and overall sequence length of PASK across these two species (Figure 5C-E). By leveraging this quaternary structural similarity, we experimentally validated the interaction between PAS-A and kinase domains (Figure 5G-H). Intriguingly, mutations in the PAC domain that disrupted PAS-C assembly (Figure 4G) also inhibited the interaction between PAS-A and the kinase domains (Figure 6I), underscoring the critical role of PAS-C domain assembly in regulating the kinase domain activity.

### The Evolutionarily Conserved PAS-C Domain of PASK likely Originated from Bacterial Oxygen Sensors

The PAS-C domain exhibits a split architecture, with its PAS fold and PAC motif originating from distant regions of the primary sequence (Figures 2A and 3B). Such a configuration of the PAS-C domain raises intriguing questions regarding its evolutionary origin. Notably, the relative length and location of the N-terminal PAS fold and C-terminal PAC motif in the primary sequence of PASK is conserved across species (Figures S7A-B). Furtherore, similar to many bacterial PAS domains, the PAS-C domain of Spotted Gars and Milkfish contains heme co-ordinating histidine in the Fα helix of the PAS-C domain. Thus, the PAS-C domain of PASK was likely acquired through horizontal gene transfer from bacteria early in metazoan evolution and underwent metazoa specific domain rearrangment, and that the ancestral PAS-C domain may have retained some capability as a heme-based oxygen sensor. However, with the water-to-land transition in the tetrapod lineage, this histidine residue was replaced by aspartate (Figure S8), potentially resulting in the loss of heme-binding capacity. Although the functional significance of this evolutionary shift remains to be elucidated, we speculate that it may have facilitated the adaptation of PASK to terrestrial environments, where nutrient sensing became more critical than oxygen sensing.

### Potential for the transition from a Partially Unfolded State to a Stronger PAS-C Domain Assembly and its impact on quaternary structure organization and catalysis

The regulatory mechanism of PASK activity is of significant interest given its essential role in stem cell differentiation and organismal metabolism. Our previous research indicated that PASK activation in response to serum stimulation involves mTORC1-mediated multisite phosphorylation, targeting sites adjacent to the PAC motif of the PAS-C domain. Remarkably, serum stimulation led to an mTORC1-dependent increase in PAS-C domain assembly (Figure 6G), suggesting that such stimulation may activate PASK by fostering quaternary structural rearrangement conducive to the interaction between the PAS-A and kinase domains. This dynamic nature of the PASK quaternary structure, modulated by the serum-responsive PAS-C domain assembly, points to a sophisticated mechanism that stabilizes the catalytic core for full activation of PASK (Figure 6H-7B).

PAS domains are known to undergo shape and fold shifts towards a partially unfolded state under physiological conditions, in response to ligand binding, or due to environmental, mechanical, or chemical signaling cues [28–31]. Because of the mode of assembly, our results suggest that the PAS-C domain of PASK might be partially unfolded under basal conditions, which undergo a signal-induced conformational change for stronger assembly of the PAS-C domain, consistent with its evolutionary origin from the bacterial PAS domain.

The discovery of a novel intramolecular folding mechanism for the PAS-C domain in PASK raises several avenues for future investigation. The precise role of the PAS-C domain in regulating PASK activity remains to be fully elucidated. While our study provides compelling evidence for the involvement of the PAS-C domain in stabilizing the catalytic core and facilitating mTORC1-mediated activation, further research is needed to determine the molecular details of this regulation. Additionally, it would be interesting to explore whether other signaling pathways or metabolites can modulate the assembly and function of the PAS-C domain.

In conclusion, our study unveils a previously unrecognized mechanism of PAS domain assembly in metazoan PASK, highlighting the dynamic and regulatable nature of protein folding. The identification of a split PAS-C domain assembled through long-range intramolecular interactions between PAS and PAC folds, along with its role in catalytic core stabilization and signal-mediated activation, represents a significant advancement in our understanding of PAS domain biology. These findings have broader implications for the study of PAS domain-containing proteins and their regulation, opening up exciting avenues for future research in this field.

## Methods and Material

### Domain Mapping and Analysis

To ascertain the domain composition, we initially utilized InterPro to simultaneously search the Conserved Domain Database (CDD), PFAM, Prosite, and CATH databases for the domain architecture of mammalian PASK [15, 16, 32, 33]. The human PASK protein sequence was also searched in the HomoloGene database of NCBI to confirm the domain architecture pattern obtained using InterPro. The human PASK sequence was analyzed separately against the SMART database [17] for domain architecture patterns. Utilizing the genomic mode of SMART, we mined the metazoan reference proteome to identify PASK orthologs that contain the domain organization order matching human PASK, while allowing for additional domains in the primary sequence. Metazoan FASTA sequences from SMART, matching the human PASK domain distribution pattern (∼297 in SMART Database), were aligned with human PASK via MAFFT 7.2 [34], followed by Guidance2 [35] scoring to compute residue-wise-confidence scores. This analysis identified PASK orthologs in the metazoan PASK, consisting of domain architecture patterns similar to human PASK. As described below, we extracted the domain architecture patterns for metazoan PASK from the SMART datasets, which were used to generate the phylogenetic tree with the PASK domain architecture datasets.

### Phylogenetic Tree Construction and Analysis

Full-length amino acid sequences for metazoans PASK were obtained from OrthoDB [36], SMART [17], and the NCBI HomoloGene database to maximize coverage of metazoans PASK. These sequences were combined and manually curated based on the length of the residues as follows: assuming the presence of at least one PAS domain (∼100 amino acids) and one kinase domain (∼250 amino acids), no metazoan sequences less than 300 amino acids were included in the final alignment. Furthermore, in species with genome duplication, only one copy of PASK was retained. After sequence curation, we generated the final alignment for jawed vertebrate PASK using MAFFT V7.2 and the G-INS-1 option. A residue-wise confidence score was computed using Guidance2 with default options. A maximum-likelihood tree was constructed from high-scoring alignments using IQ-TREE2 [37], with the JTT+F+R5 (BIC approximation) model and 1,000 bootstrap replicates. Trees were visualized using iTOL with bootstrap values and domain distribution patterns extracted from annotated SMART [38].

### AlphaFold Model Generation and Analysis

When available, AlphaFold models were imported directly into ChimeraX(v1.7) [39]. Full-length structural models of rifleman, West African lungfish, and Himalayan leaf-nosed bats were generated using the ColabFold notebook with Amber molecular dynamic relaxation [19]. No templates were used to generate structural models, and a minimum of five models were created and analyzed using the pLDDT and PAE scores. The best-scoring models were further analyzed for comparative structural phylogenetic analyses.

### Model Validation Workflow

We constructed structural models of the individual domains, PAS-B and PAS-C, using RosettaFold and ESMFold [21, 22]. Residues corresponding to the PAS-B domain were modeled using template-free, deep-learning-based RosettaFold. For the PAS-C domain, we combined residues 333-405, which included N-terminal PAS-folds, and residues 886-933, which included C-terminal PAC-folds from human PASK. Structural models were superimposed, and models with the highest average pLDDT score and lowest Angstrom error were selected for further analysis. Similarly, ESMFold structures were generated using the ColabFold notebook from PAS-A and PAS-C sequences. The resulting PDB files from RosettaFold or ESMFolds were searched against the PDB database using FoldSeek [20]. The highest-scoring hits from the FoldSeek output were structurally aligned in ChimeraX using MatchMaker. PDB files from RosettaFold and ESMFolds were also searched against the CATH structural family database, and aligned sequences from the CATH Functional Family were generated to identify and build a structure-based sequence alignment with the human PAS-C domain.

### Construction of improved HMMER3 Profile and Structure-Based Sequence Alignment

PAC domain sequences were extracted from the SMART database (accession number: SM00086), site motif (Prosite matrix, accession number: PRU00141), and the PDB database for PAS domain structures from eukaryotic and prokaryotic proteins. These sequences were combined with the PAC domain sequences from 19 invertebrate and vertebrate PASK species, as identified by SMART (Figure S4). An MSA was generated using MAFFT V7.2 using the G-INS-I (slowest, most accurate) approach, which was used to build an HMMER3 profile using the Unipro UGENE software. Hmmsearch was performed against the metazoan reference proteome. Structure-based sequence alignments were generated from MSA and structural templates using the ES Pript 3.0 [40].

### Plasmids for Expression of Human PASK and Cloning of Rifleman PASK

Plasmids for the expression of human PASK cDNA have been previously described [24]. All mutations were made using a Q5 mutagenesis kit (New England Biolabs), according to the manufacturer’s protocol. cDNA corresponding to rifleman PASK was synthesized and cloned into the pQCXIP-GFP vector. The published rifleman PASK sequence lacks the first 50 residues of its primary sequence. Therefore, we added missing amino acids from the full-length sequence of zebra finch PASK, which shares more than 90% sequence identity with rifleman PASK, excluding the gaps caused by PAS-C domain splitting.

### Cell Lines and Myoblast Culture and Differentiation

HEK293T and C2C12 cells were maintained in Dulbecco’s Modified Eagle’s medium (high glucose) supplemented with 10% Fetal Bovine Serum and 1% penicillin/streptomycin. HEK293T cells were transfected using a Polyethylenimine (PEI)-based transfection reagent. Where the effect of serum was used, cells were serum starved overnight in DMEM + 1% PS. The cells were stimulated with 20% serum for 20 min or 2 h before cell lysis and downstream applications. To evaluate the functional orthology of rifleman PASK in the myogenesis assay, we silenced C2C12 cells with Smartpool siRNA (Dharmacon) against mouse PASK using Lipofectamine. 24 hours after siRNA transfection, C2C12 cells were infected with a purified retrovirus carrying GFP (as a control) or rifleman-GFP. Cells were allowed to reach confluency (for an additional 24 h) before stimulating myogenesis by replacing the growth medium with media containing 2% horse serum for 48 h. At the end of the differentiation time course, the cells were fixed and myogenesis was visualized using an MF20 antibody (anti-MHC). The fusion index was calculated as the number of nuclei inside MHC + myofibers divided by the total number of nuclei per field. Ten independent fields were examined in total.

### Cell-lysis, Co-Immunoprecipitation, and Western Blotting

For co-immunoprecipitation experiments, HEK293T cells were lysed in native lysis buffer as described previously. Bait proteins (V5- or Flag-tagged) were immunoprecipitated from clarified cell lysates using anti-V5 or anti-Flag magnetic beads. The beads were washed vigorously using NTB wash buffer (50 mM Tris, pH 7.6, 150 mM NaCl, and 1% Triton-X100). The immunoprecipitants were denatured using Laemmli buffer. Proteins were separated by SDS-PAGE, and western blotting was performed using the indicated antibodies.

For the TEV protease cleavage system, we inserted the Flag-TEV-V5 cassette internally in human PASK between amino acids 735-738. HEK293T cells expressing WT or PAC-mutated Human PASKFlag-TEV-V5 were lysed, and FLAG-tagged PASK was purified from clarified cell extracts using anti-Flag magnetic beads. Flag beads were washed four times with a relatively high salt buffer (250mM NaCl, 0.03% sodium deoxycholate, 1% Triton X-100 in Tris HCl, pH 7.6 buffer) to remove any interacting proteins that may bridge the N-terminal and C-terminal fragments, as well as dissociate any protein dimers that may be present in PASK. Beads were then washed with normal salt IP wash buffer (150mM NaCl, 1% Trixon-100 in Tris HCl, pH 7.6 buffer) and once with TEV cleavage buffer (50mM Tris HCl, pH 8.0, 0.5mM EDTA, 1mM DTT). Proteins were eluted using Flag-peptides in TEV protease cleavage buffer. Part of samples were saved as input control for uncleaved sample. Flag eluents were cleaved with TEV protease at 25C for 4 h on a nutator, and samples were saved as input controls for cleaved samples. The cleaved samples were recaptured using anti-V5 antibodies overnight or for 4 hours at 4OC. Immunoprecipitants were washed with IP wash buffer and proteins were released from the beads using Laemmli buffer. Proteins were separated on SDS-PAGE gels, and Flag and V5 tagged proteins were detected using the indicated antibodies.

### In Situ Cysteine Cross-linking Analysis (VivoX)

We adapted the VivoX method from a recent publication [25] to determine the interdomain communication in this study and extended it to study the signal-regulated quaternary structure reorganization of PASK. HEK293T cells expressing various combinations of WT and cysteine-mutated PASK were treated with 180µM 4-DPS or vehicle (DMSO) for 20 min at 37°C. Since PASK is a soluble cytoplasmic protein, we modified the cell lysis buffer used for VivoX. HEK293T cells were lysed in modified RIPA lysis buffer (50mM Tris-HCl, pH 7.6, 1% NP-40, 1% Sodium Deoxycholate, 0.2% SDS) containing 50mM N-ethylmaleimide (NEM), 1X protease inhibitor, and phosphatase inhibitor cocktail. The cell extracts were clarified by centrifugation at 4°C for 20 min at 15,000 rpm. Clarified cell extracts were denatured using SDS-PAGE Laemmli buffer with (reducing) or without 5% β-ME (non-reducing). SDS-PAGE and western blotting were performed as previously described. To determine the effect of serum on the quaternary structure using VivoX, HEK293T cells were serum-starved overnight in a serum-free medium containing DMEM + 1% PS. The next day, the cells were treated with 50µM 4-DPS or vehicle (DMSO) with or without 20% serum for 20 min. Cell lysis and western blotting were performed as previously described.

## Author Contributions

CKK conceived the project and performed phylogenetic analysis, computational structure modeling, and sequence analysis. SD, MX, and RPP performed experiments and data analysis. C. K. K. wrote the manuscript with input from all authors.

## Supporting information

HMMER profile Dataset Files

## Acknowledgments

We are grateful for the lively discussion and helpful suggestions regarding the phylogenetic components of this paper with colleagues Drs. Jeramiah Smith, Phil Skipwith, Rosana Zenil Ferguson, and members of Kikani lab. We thank Dr. Phil Skipwith for pointing out the ancestral relationship between rifleman and passerine birds, which informed the domain mapping and structure superposition. This work was supported by funding from the National Institute of Health (R01 AR073906 to C. K.).

## Associated Data Files

The improved HMMER3 profile file for PAC domain prediction is available in Supplementary Data File 1.

