## Supplemental document and Figures for "Nutrient Signaling-Dependent Quaternary Structure Remodeling Drives the Catalytic Activation of metazoan PASK"

### Nutrient Sensing and Catalytic Activation through Quaternary Structure Remodeling in the Metazoan PASK

Supplementary Figure Legends

Figs. S1-S10

Movies S1

Data S1

**Figure S1: Mammalian PASK Gene Tree with Domain Architecture Analysis**

Mammalian PASK sequences were extracted from UniProt, OrthoDB, or NCBI, and a multiple sequence alignment was generated using MAFFT. A phylogenetic tree was constructed using IQ-Tree2 and the resulting Newick file was visualized using iTOL. The domain architecture of the corresponding mammalian PASK was obtained using SMART and visualized using iTOL.

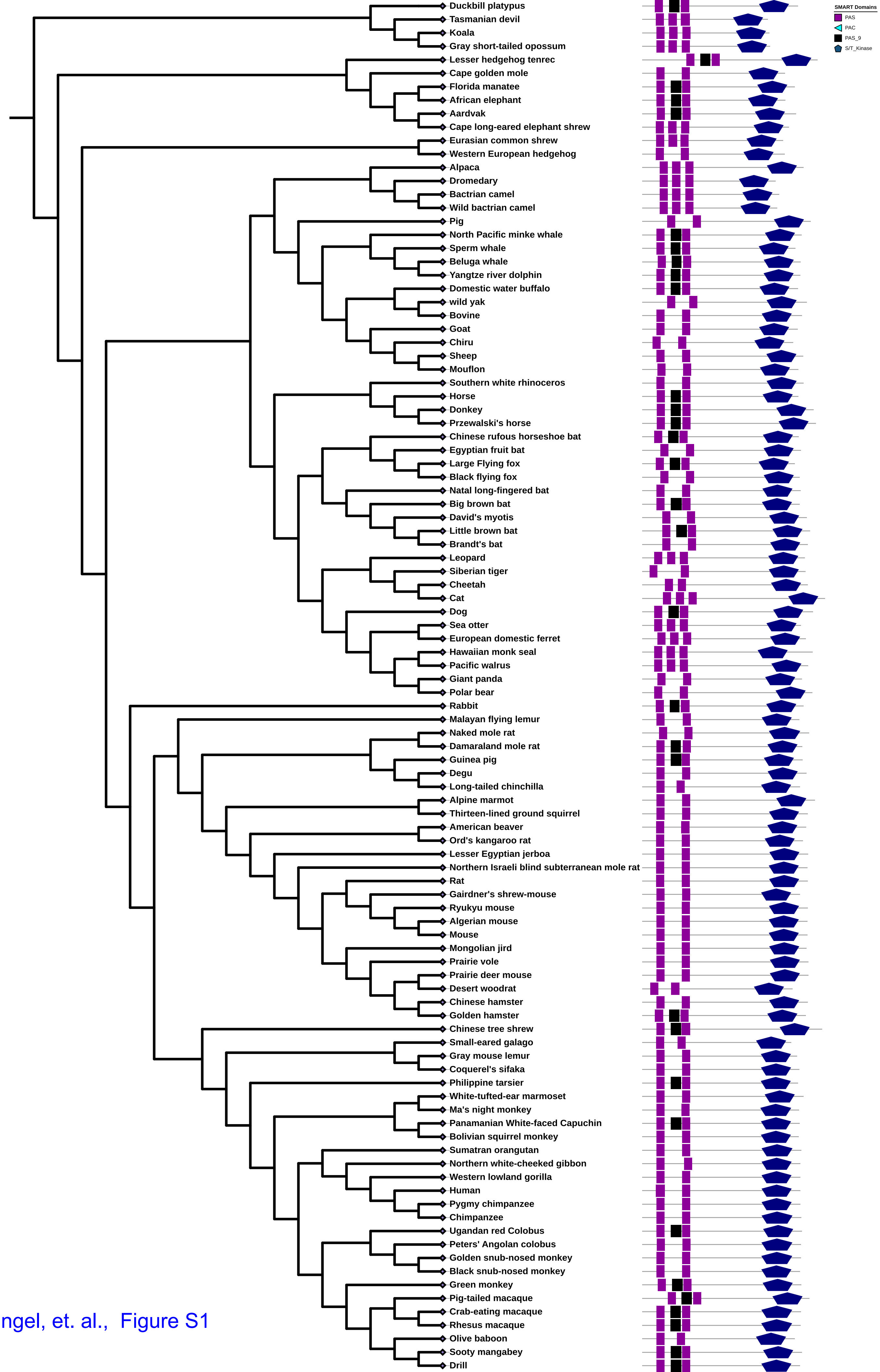


Dhungel, et. al., Figure S1

**Figure S2: Canonical PAS Domain Structure from Bacterial FixL**

Secondary structural elements corresponding to the PAS-fold-(purple) and PAC-fold (cyan) are depicted in the bacterial FixL crystal structure (PDB: 1DP6) with a coordinated heme. The structure-based alignment illustrates the position of conserved residues across the PAS and PAC folds in bacterial, human, and mouse PAS domains, highlighting a significant diversity in the primary sequence that folds into a conserved PAS domain.

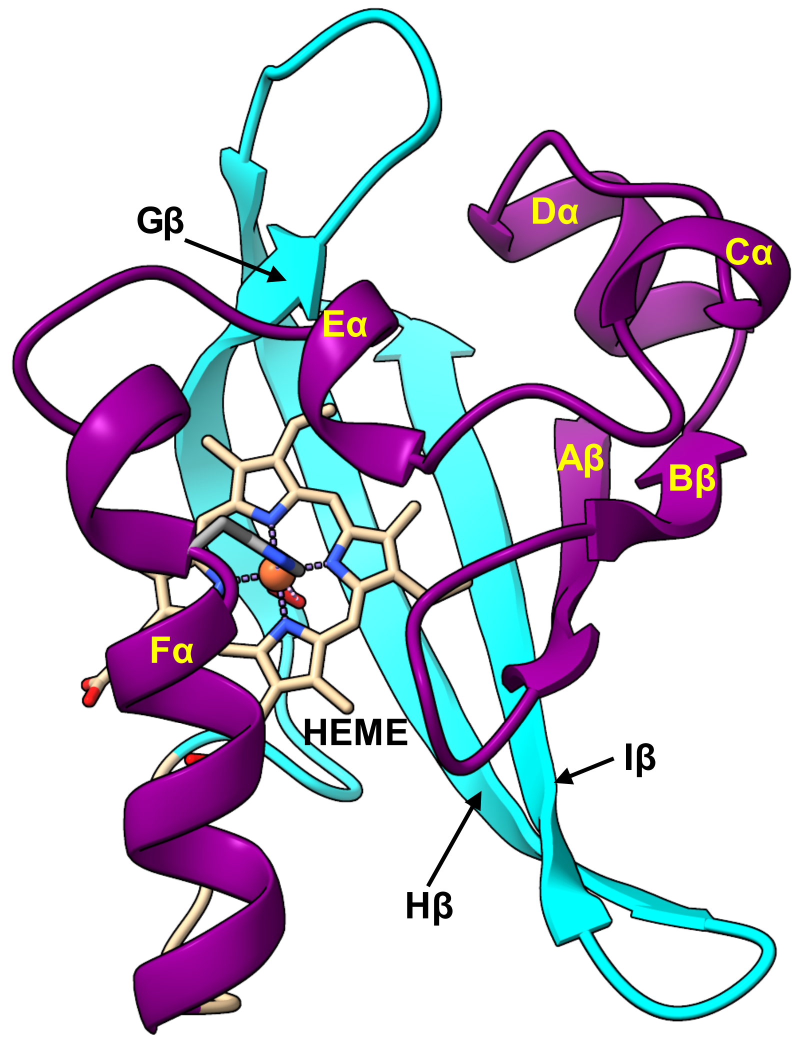


**PAS Fold**

```
FIXL_BJAPO                               Aβ       TT   Bβ   Cα        Dα          Eα           Fα
                                             20            30          40          50          60          70
FIXL_BJAPO    ...........TIPD...........AMIVID...GHGIIQLFSTAAERLFGWSELEAIGQN..VNILMPEPDRSRHDSYISRYRTTSDPH.
DOSP_ECOLI    .............PA.....LEQNMMGAVLIN...ENDEVMFFNPAAEKLWGYKREEVIGNN..IDMLIPRDLRPAHPEYIRHNREGGKAR.
NIFL_AZOVI    ............API...........AISITD...LKANILYANRAFRTITGYGSEEVLGKN..ESILSNGTTPRLVYQALWGRLAQKKP..
PHOT_BACSU    .......................VRVGVVITDPALEDNPIVYVNQGFVQMTGYETEEILGKN..CRFLQGKHTDPAEVDNIRTALQNKEP..
KCNH2_HUMAN   .......................QSRKFIIANARVENCAVIYCNDGFCELCGYSRAEVMQRPCTCDFLHGPRTQRRAAAQIAQALLGAEE..
ARNT_HUMAN    ..........................PTEFISRHNIEGIFTFVDHRCVATVGYQPQELLGKN.IVEFCHPEDQQLLRDSFQQVVKLKGQV..
CLOCK_MOUSE   LVAIGRLHS.HMVPQPANGEIRVK..SMEYVSRHAIDGKFVFVDQRATAILAYLPQELLGTS.CYEYFHQDDIGHLAECHRQVLQTREKI..
EPAS1_HUMAN   ..........................SKTFLSRHSMDMKFTYCDDRITELIGYHPEELLGRS.AYEFYHALDSENMTKSHQNLCT.KGQV..
BMAL1_MOUSE   FVATVRLAT....PQFIKEMCTVEEPNEEFTSRHSLEWKFLFLDHRAPPIIGYLPFEVLGTS.GYDYYHVDDLENLAKCHEHLMQ.YGKG..
PER_DROME     VICATPIKSSYKVPD...EILSQK..SPKFAIRHTATGIISHVDSAAVSALGYLPQDLIGRS.IMDFYHHEDLSVMKETYETVMK.KGQT..
Per_Mouse     LLLAERVHSGYEAPR.....IPPE..KRIFTTTHTPNCLFQAVDERAVPLLGYLPQDLIETP.VLVQLHPSDRPLMLAIHKKILQAGGQP..
PYP_HALHA     ..........................GAIQLD...GDGNILQYNAAEGDITGRDPKQVIGKNFFKDVAPCTDSPEFYGKFKEGVASGNLNT
```

**PAC Fold**

```
FIXL_BJAPO         TT     Gβ      TT        Hβ       ..  TT     Iβ
                      80         90         100        110        120        130
FIXL_BJAPO    ..IIGIGRIVTGKRRDGTTFPMHLSIGEM..QSGGEPYFTGFVRDLTEHQQTQARLQELQ................................
DOSP_ECOLI    ..VEGMSRELQLEKKDGSKIWTRFALSKV..SAEGKVYYLALVRDASVEMAQKEQTRQLI................................
NIFL_AZOVI    .....WSGVLVNRRKDKTLYLAELTVAPVLNEAGETIYYLGMHRDTSE............................................
PHOT_BACSU    .....VTVQIQNYKKDGTMFWNELNIDPM..EIEDKTYFVGIQNDITEHQQTQARLQELQSELVHVSRLSAMGEMASALAHELNQPLAAISN
KCNH2_HUMAN   .....RKVEIAFYRKDGSCFLCLVDVVPVKNEDGAVIMFILNFEVVMEK...........................................
ARNT_HUMAN    .....LSVMFRFRSKNQEWLWMRTSSFTFQNPYSDEIEYIICTNTNVKNSSQE.......................................
CLOCK_MOUSE   .....TTNCYKFKIKDGSFITLRSRWFSFMNPWTKEVEYIVSTNTVVLANVLE.......................................
EPAS1_HUMAN   .....VSGQYRMLAKHGGYVWLETQGTVIYNPRNLQPQCIMCVNYVLSEIEKN.......................................
BMAL1_MOUSE   .....KSCYYRFLTKGQQWIWLQTHYYITYHQWNSRPEFIVCTHTVVSYAEVRAE.....................................
PER_DROME     AGASFCSKPYRFLIQNGCYVLLETEWTSFVNPWSRKLEFVVGHHRVFQGPKQCNVFEAAP................................
Per_Mouse     ....FDYSPIRFRTRNGEYITLDTSWSSFINPWSRKISFIIGRHKVRVGPLNEDVFAAPP................................
PYP_HALHA     YQMTPTKVKVHMKK.................ALSGDSYWVFVKRV...............................................
```

Dhungel, et. al., Figure S2

**Figure S3: Predicted Domain Architecture of Human PAS Domain-Containing Proteins Shows Inconsistency in Identifying Tandem PAS and PAC Folds**

**A.** Protein sequences corresponding to PAS domain-containing proteins were extracted from UniProt and aligned using MAFFT. The human PAS domain family tree was generated using the IQ-Tree2. The protein domain architecture was extracted from SMART and visualized with the protein family tree from IQ-Tree2 using iTOL.

**B.** Experimentally derived structures of the ARNT-PAS-A domain (PDB: 5Sy5) and PAS-B domain (PDB: 2b02). The domain architecture was annotated above the structural models. Note that the PAS-A domain boundary extends beyond the SMART-predicted domain boundary and the accurate prediction of PAS and PAC folds for the PAS-B domain by SMART.

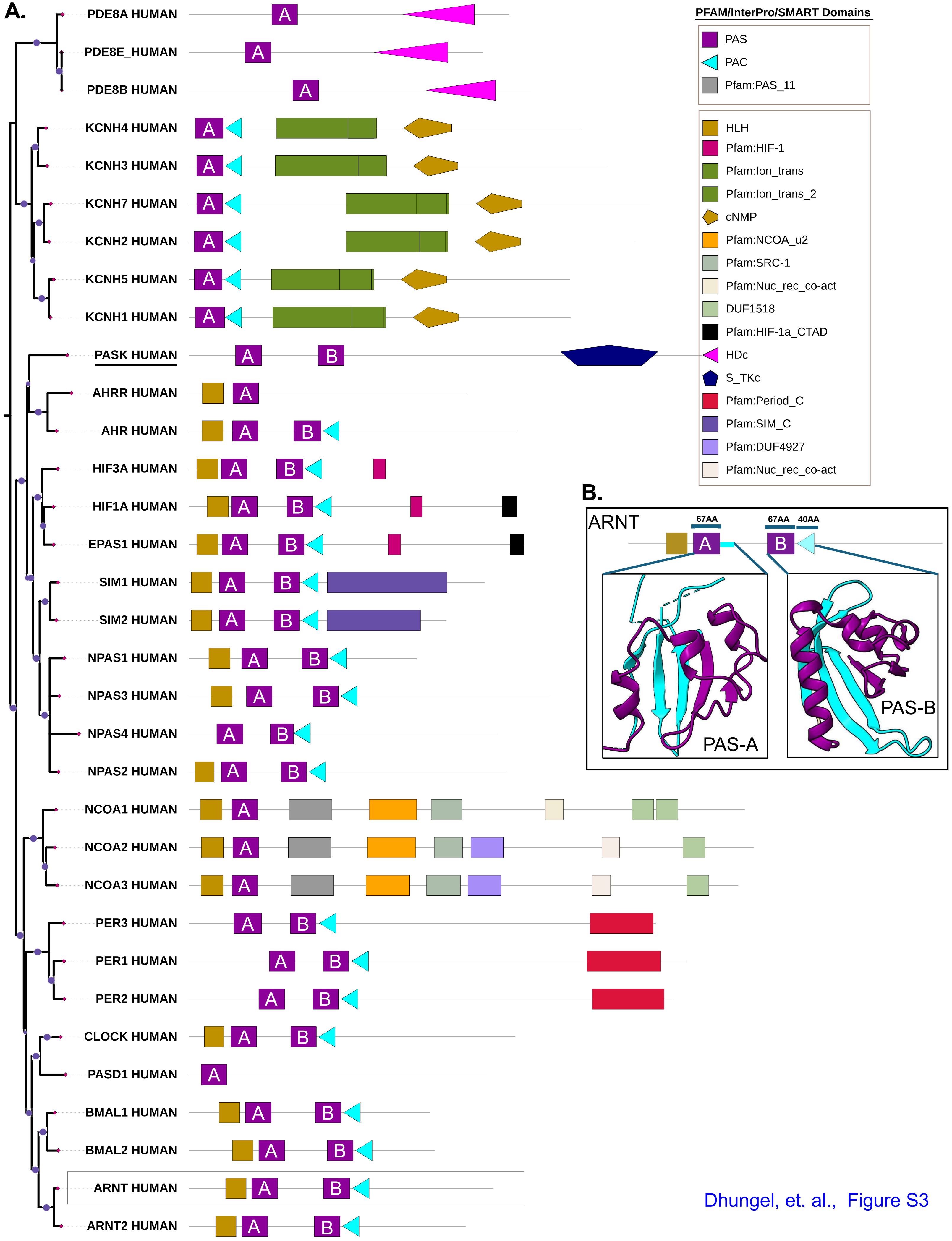


Dhungel, et. al., Figure S3

**Figure S4: Metazoan PASK with Predicted PAC Domains**

A metazoan-wide PASK protein domain tree identified 19 species, for which SMART/HMMER predicted the presence of a single PAC domain. The domain architecture of PASK in invertebrate and vertebrate species is shown.

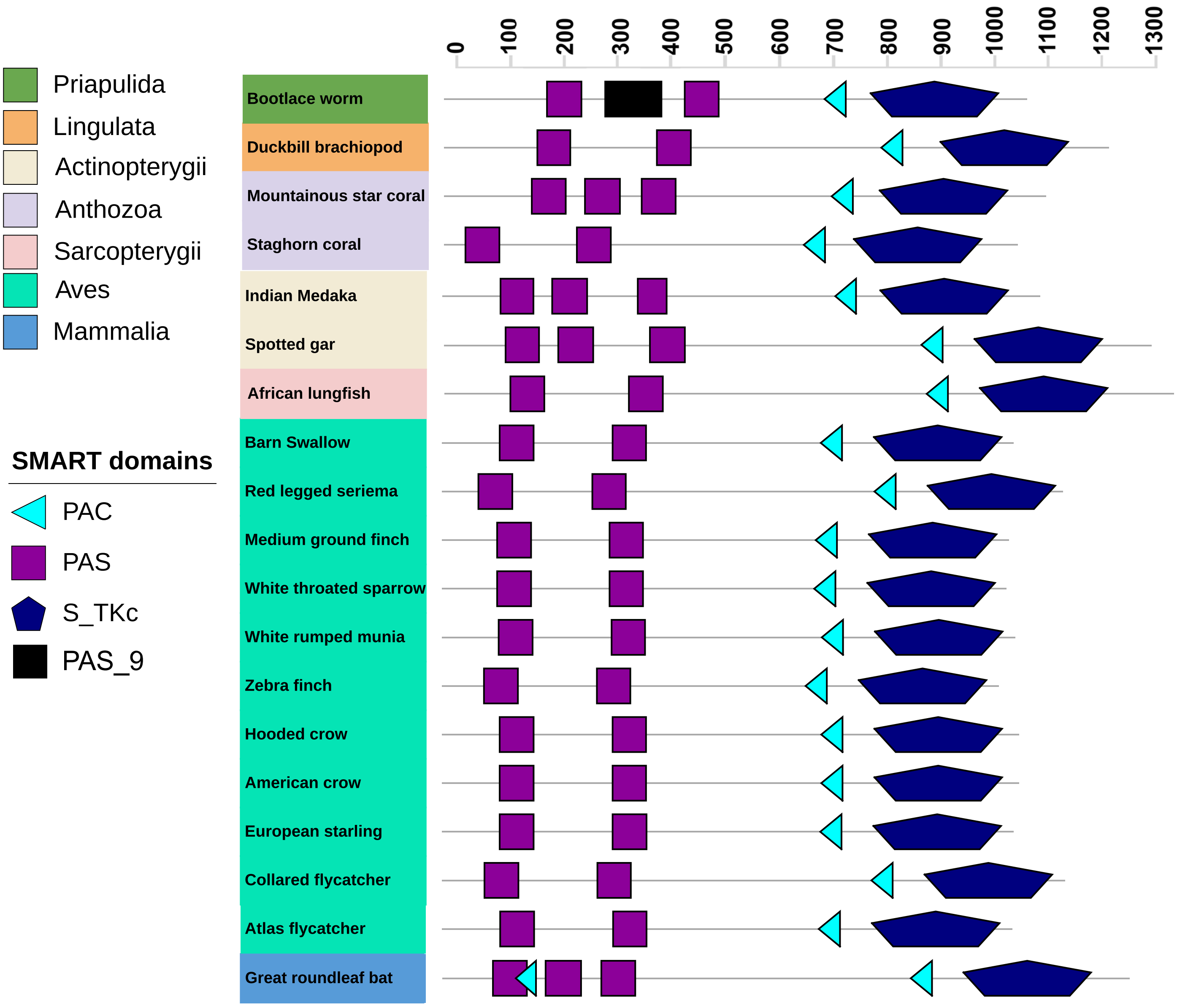


Dhungel, et. al., Figure S4

**Figure S5: Sequence-Structural Conservation of PAS-A and PAS-B Domains of PASK Across Selected Jawed Vertebrates**

**A.** AlphaFold models of PAS-A domains of PASK from selected vertebrates showing structural similarity and a high level of sequence identity.

**B.** AlphaFold models of the PAS-B domains of PASK from selected vertebrates. Note that the rifleman PAS-B domain is not well modeled in the AlphaFold structure, possibly because of missing key residues in the current version of the genome assembly.

A.

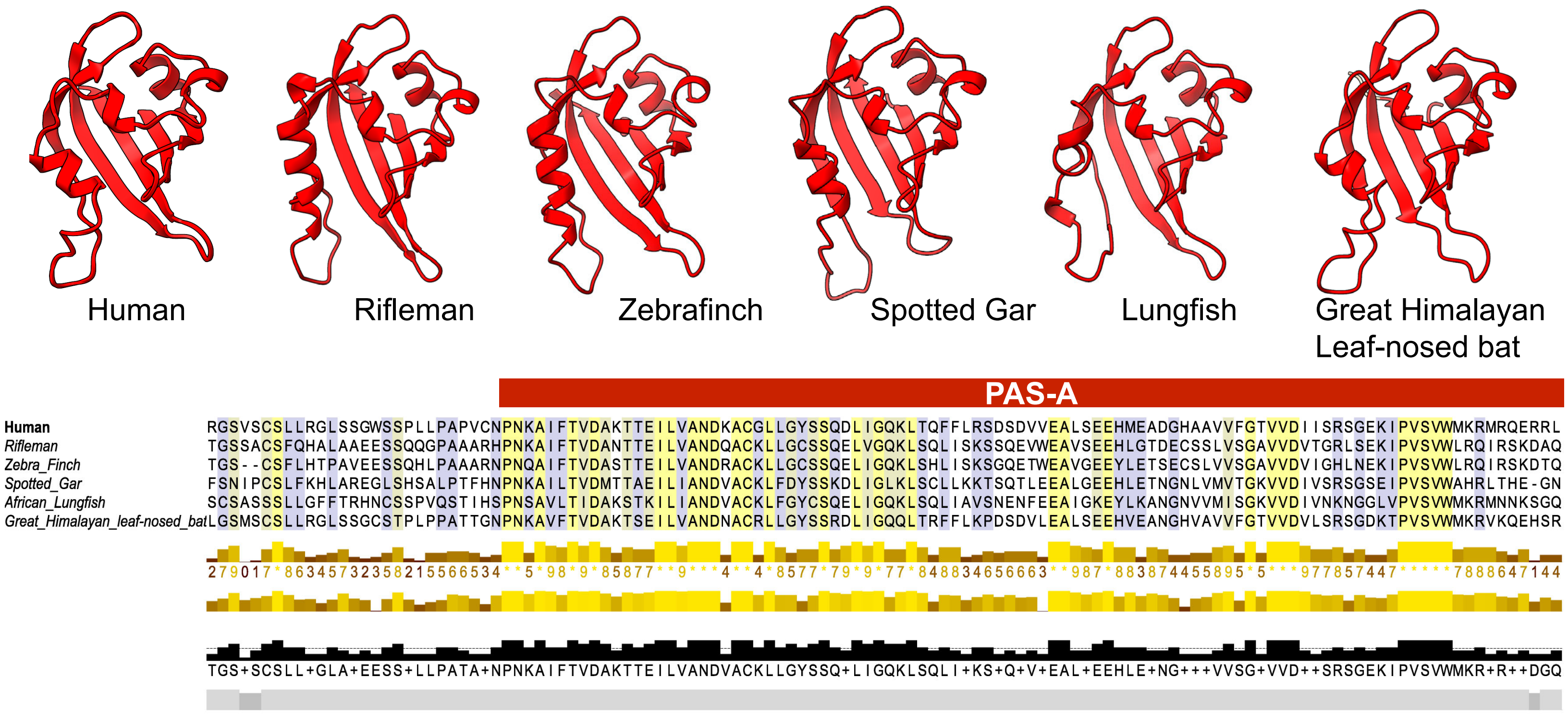


B.

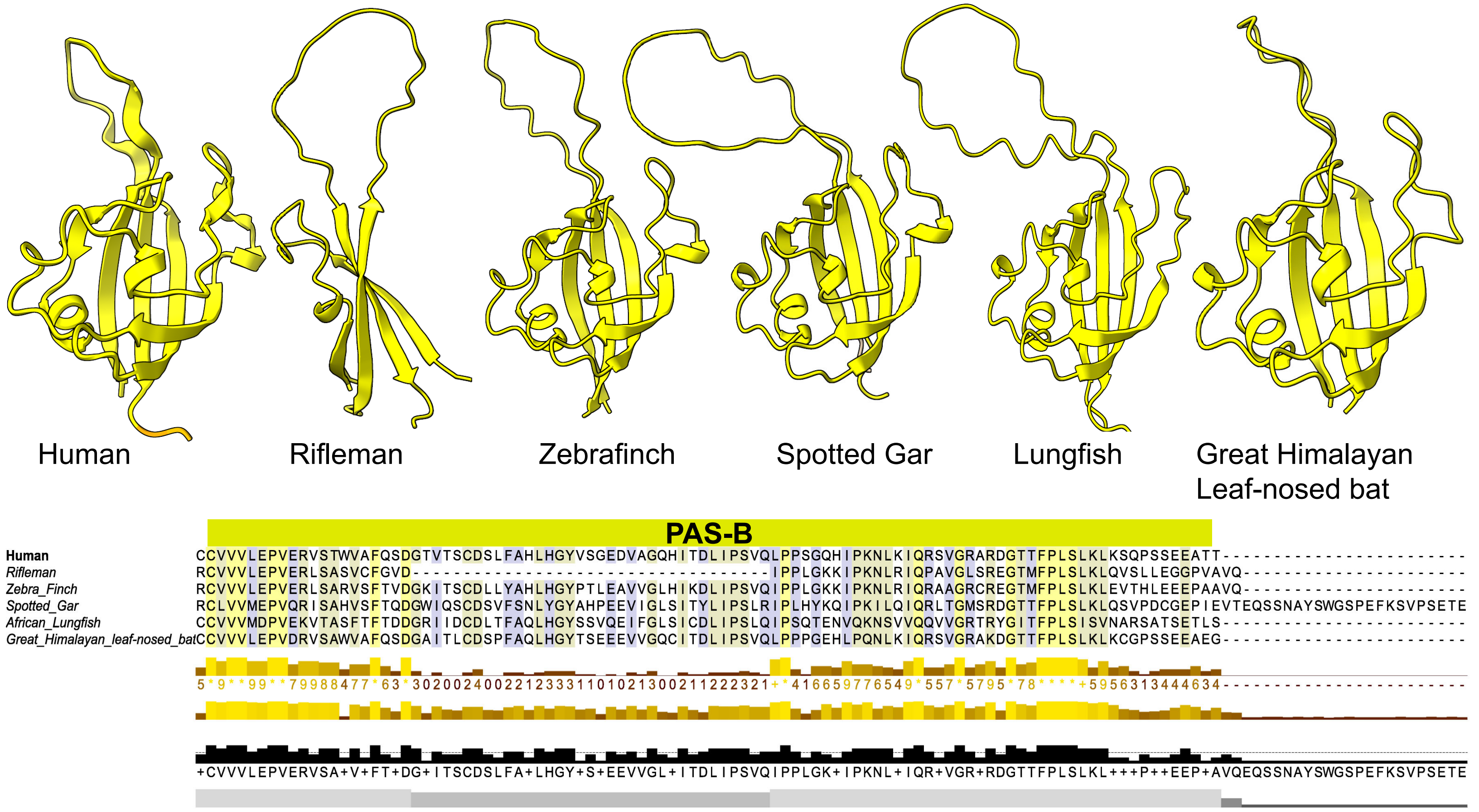


Dhungel, et. al., Figure S5

**Figure S6: Development and Validation of i-HMMER3 Profile to Identify PAC Motif in Large-Scale Proteome Sequence**

**A.** The flowchart describes the source of the PAC domain sequences, MSA construction, and generation of the HMMER3 profile from MSA. Please see the Methods section for further details.

**B.** Logo output from the improved HMMER3 (i-HMMER3) profile.

**C.** Number of PAS domain-containing proteins identified by SMART, Prosite, or i-HMMER3 in the reference human proteome.

**D.** Comparison of the performance of the HMMER3 profile in SMART vs. i-HMMER3 in identifying PAS domain-containing proteins across metazoa. The HMMER profile from SMART or i-HMMER3 was used to search for the metazoan reference proteome using the hmmsearch function of HMMER.

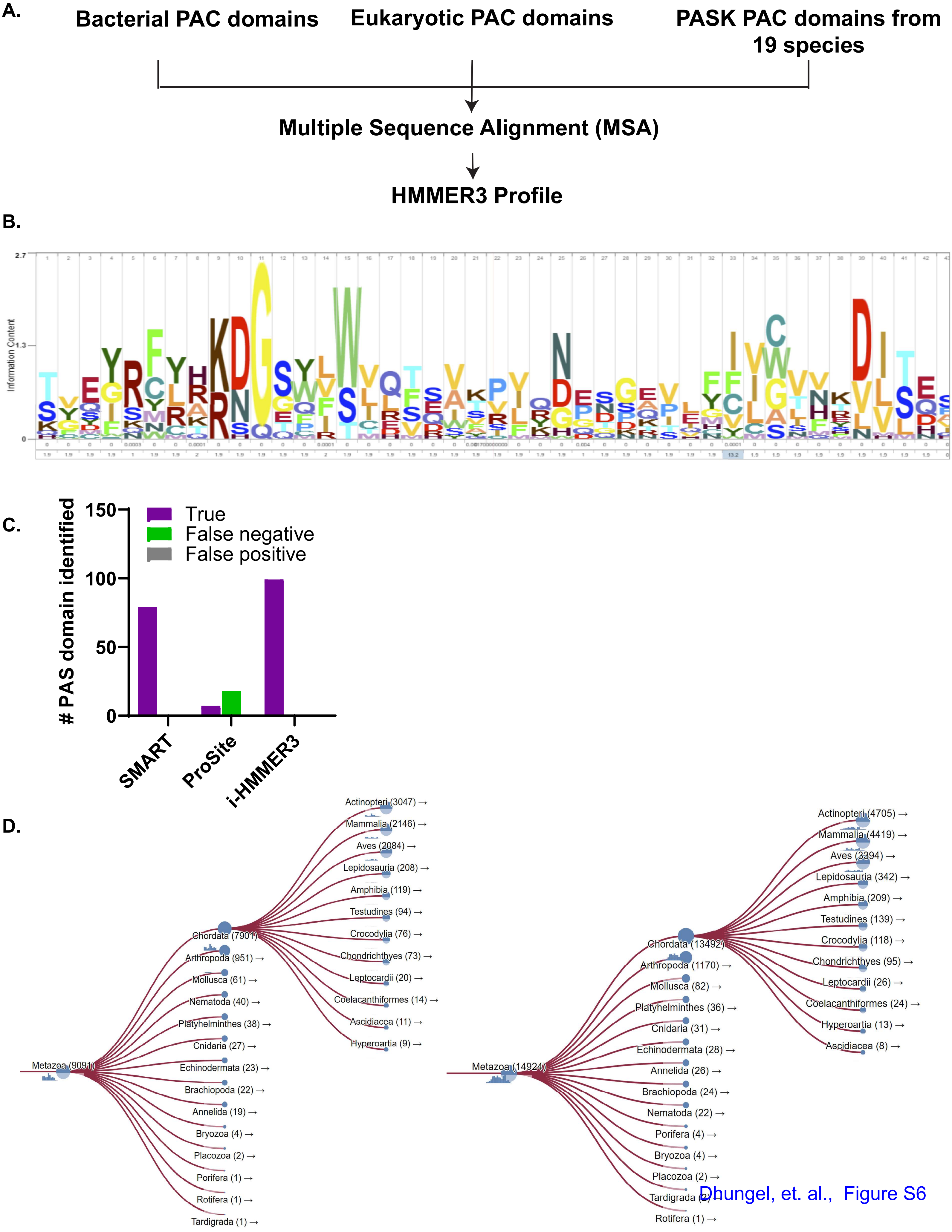


Dhungel, et. al., Figure S6

**Figure S7: Identification of PAC Motif in Metazoan PASK**

**A.** The metazoan reference proteome was searched for PAC motifs using i-HMMER3. The resulting hits were filtered for PASK, and domain pattern information was extracted from the i-HMMER3 output and converted into an iTOL-compatible dataset domain file. The domain architecture was visualized using iTOL for the corresponding jawed vertebrate PASK.

**B.** Structural elements that encompass the PAS-C domain, including the PAS-fold (purple), PAC-fold (cyan), and a 486 amino acid long loop connecting the two folds. The PAS-C domain is connected to the kinase domain via a 68 amino acid long loop consisting of a single alpha-helix and an unstructured loop.

**C.** Model depicting assembly of the PAS-C domain in PASK across jawed vertebrates.

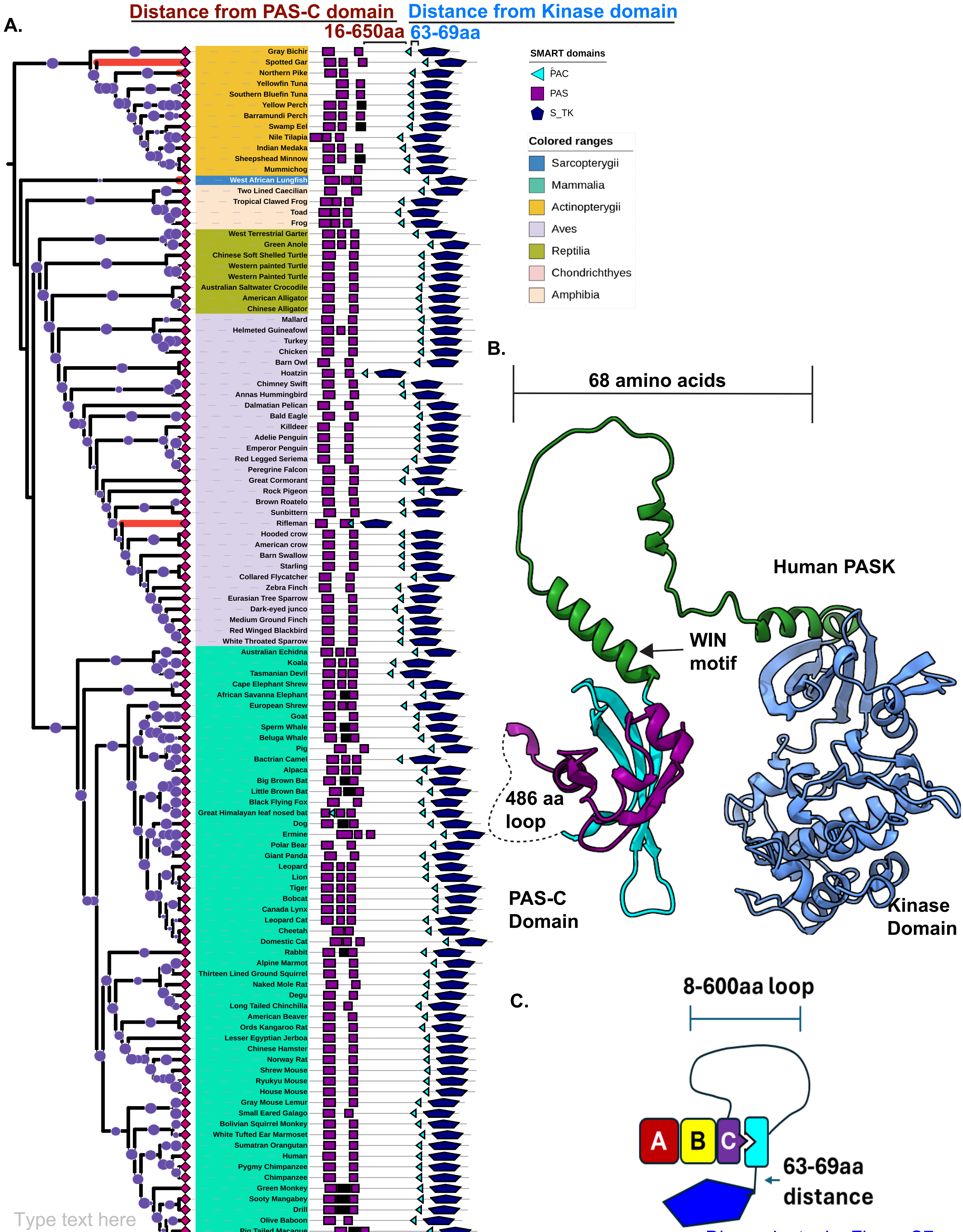


Dhungel, et. al., Figure S7

**Figure S8: Structure-Based Sequence Alignment of PAS-C Domain of PASK with Bacterial Heme-Coordinating PAS Domain**

PAS-C domain sequences from selected jawed vertebrates, including fish, amphibians (marked by A), reptiles (marked by R), birds, and mammals, were aligned with FixL from *Bradyrhizobium japonicum* (FixL_BRAJA), *Rhizobium meliloti* (FixL_RHIME), or *E. coli* oxygen sensor, DOSP_ECOLI using MAFFT. Structure-based alignment was generated using ESPript 3.0, with FixL_BRAJA as a template.

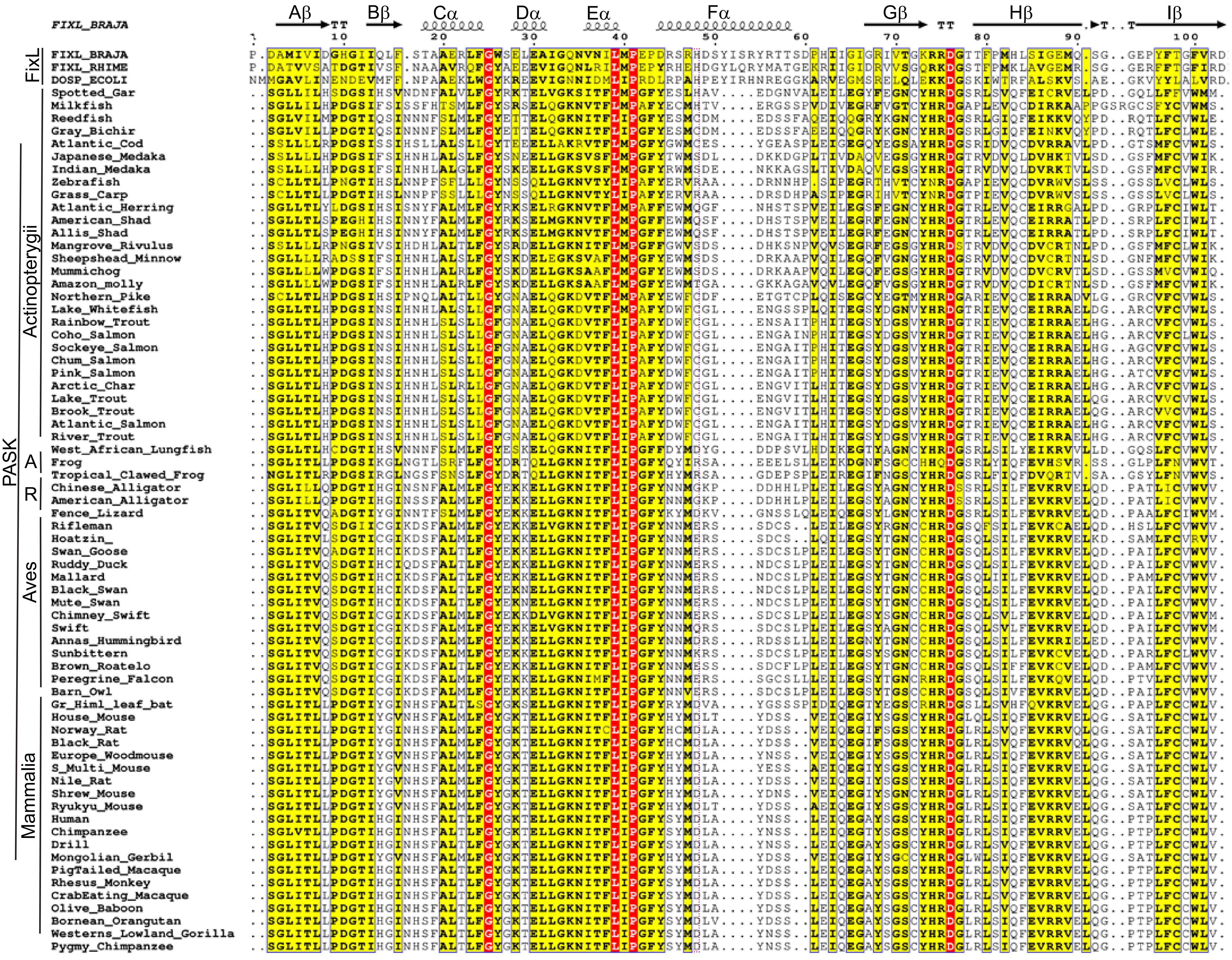

FIXL_BRAJA
Aβ
Bβ
Cα
Dα
Eα
Fα
Gβ
Hβ
Iβ
TT
1
10
20
30
40
50
60
70
80
90
100
FixL
PASK
Actinopterygii
A
R
Aves
Mammalia
FIXL_BRAJA
FIXL_RHIME
DOSP_ECOLI
Spotted_Gar
Milkfish
Reedfish
Gray_Bichir
Atlantic_Cod
Japanese_Medaka
Indian_Medaka
Zebrafish
Grass_Carp
Atlantic_Herring
American_Shad
Allis_Shad
Mangrove_Rivulus
Sheepshead_Minnow
Mummichog
Amazon_molly
Northern_Pike
Lake_Whitefish
Rainbow_Trout
Coho_Salmon
Sockeye_Salmon
Chum_Salmon
Pink_Salmon
Arctic_Char
Lake_Trout
Brook_Trout
Atlantic_Salmon
River_Trout
West_African_Lungfish
Frog
Tropical_Clawed_Frog
Chinese_Alligator
American_Alligator
Fence_Lizard
Rifleman
Hoatzin_
Swan_Goose
Ruddy_Duck
Mallard
Black_Swan
Mute_Swan
Chimney_Swift
Swift
Annas_Hummingbird
Sunbittern
Brown_Roatelo
Peregrine_Falcon
Barn_Owl
Gr_Himl_leaf_bat
House_Mouse
Norway_Rat
Black_Rat
Europe_Woodmouse
S_Multi_Mouse
Nile_Rat
Shrew_Mouse
Ryukyu_Mouse
Human
Chimpanzee
Drill
Mongolian_Gerbil
PigTailed_Macaque
Rhesus_Monkey
CrabEating_Macaque
Olive_Baboon
Bornean_Orangutan
Westerns_Lowland_Gorilla
Pygmy_Chimpanzee

**Figure S9: Structure-Based Sequence Alignment of Mammalian PAC Domain of PASK with Bacterial and Eukaryotic PAC Domains in SMART/HMMER Database**

**A.** PAC domain sequences were extracted from the SMART dataset and aligned with the human and Great Himalayan leaf-nosed bat PASK. A structure-based alignment was generated using full-length human PASK as a template, which confirmed the structural alignment of the PAS-C PAC domain with the bacterial and eukaryotic PAC domains. **B.** Consensus motif for PAC domain alignment from A.

**A.**

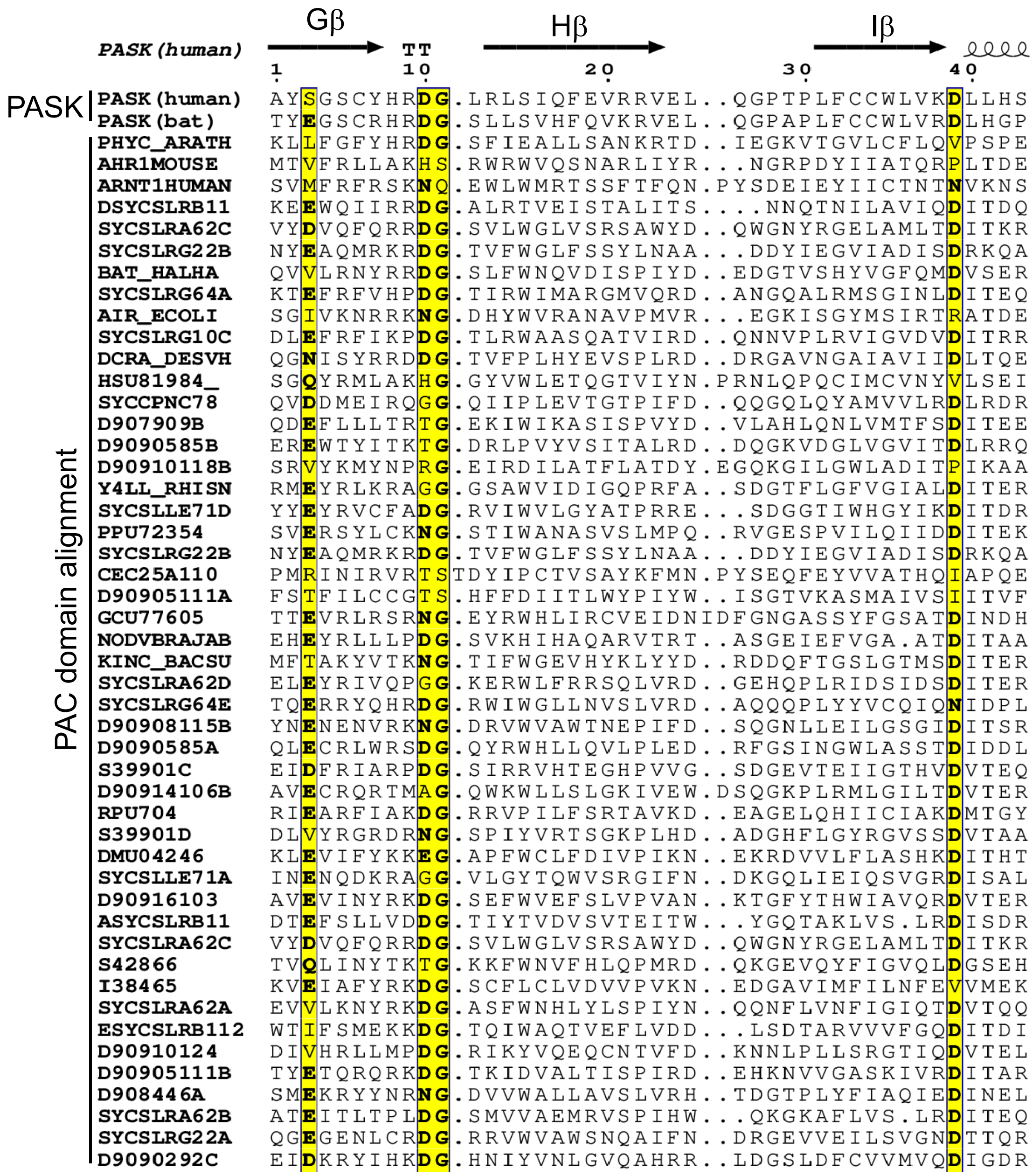


**B.**

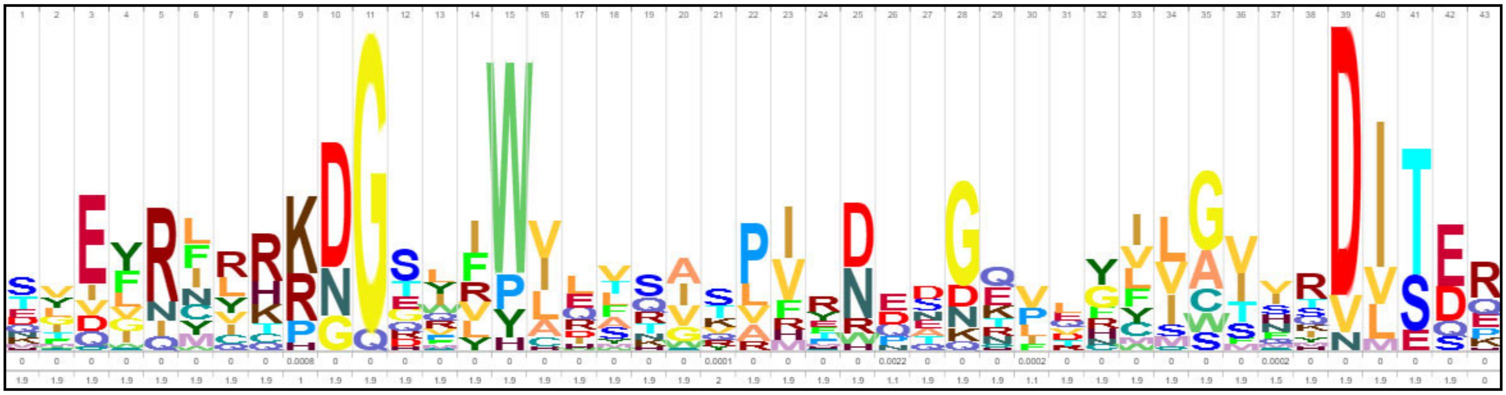

Dhungel, et. al., Figure S9

**Figure S10. Principle of PADASC workflow**

A phylogenetic gene tree was built using curated MSA constructed from diverse taxa. Domain architecture analysis and deep-learning-based structural models were analyzed at the evolutionary scale. Finally, intra-and inter-domain predicted aligned errors (PAE) were compared between quaternary structures from AlphaFold to identify consistent patterns of residue interactions across quaternary structures.

### Phylogeny Assisted Domain Architecture analysis and Structure Comparison

#### PADASC approach

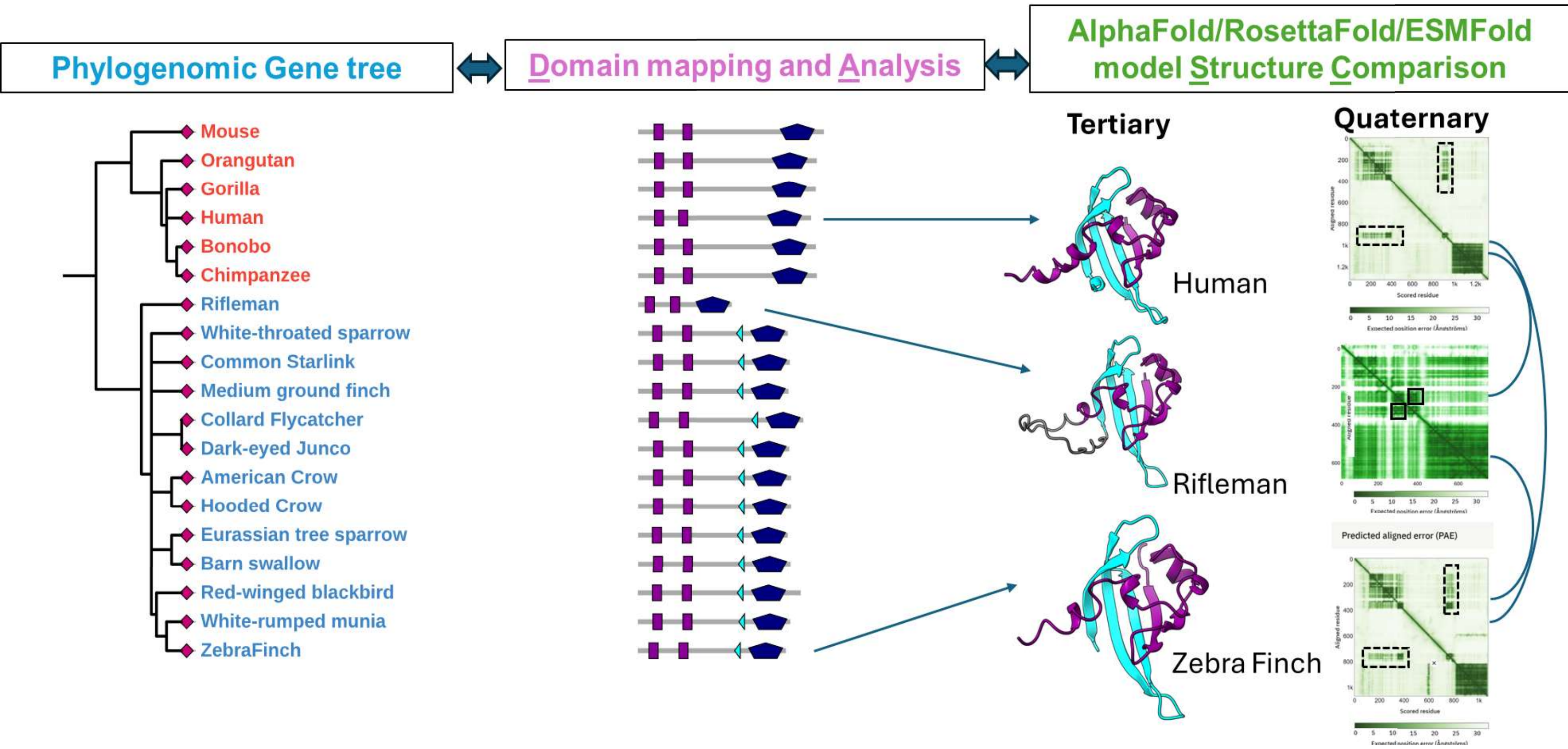


Dhungel, et. al., Figure S10
