## Supplementary material for "Nutrient Signaling-Dependent Quaternary Structure Remodeling Drives the Catalytic Activation of metazoan PASK": HMMER profile Dataset Files

```
HMMER3/f [3.3.1 | Jul 2020]
NAME  All_Human_PAC_with_hPASK_alignment
LENG  62
ALPH  amino
RF    no
MM    no
CONS  yes
CS    no
MAP   yes
DATE  Wed Feb 21 09:37:55 2024
NSEQ  213
EFFN  36.144608
CKSUM 158212377
STATS LOCAL MSV       -8.4466  0.71889
STATS LOCAL VITERBI   -8.5992  0.71889
STATS LOCAL FORWARD   -4.1886  0.71889
HMM          A        C        D        E        F        G        H        I        K        L        M
      N        P        Q        R        S        T        V        W        Y
            m->m     m->i     m->d     i->m     i->i     d->m     d->d
  COMPO   2.76119  4.12579  2.90808  2.63397  3.40029  2.91396  3.63367  2.70207  2.95647  2.41953
3.86266  3.29050  3.58042  2.89985  2.60516  2.64466  2.74060  2.60754  4.06683  3.48214
          2.68618  4.42225  2.77519  2.73123  3.46354  2.40513  3.72494  3.29354  2.67741  2.69355
4.24690  2.90347  2.73739  3.18146  2.89801  2.37887  2.77519  2.98518  4.58477  3.61503
          0.39046  4.80226  1.15504  0.61958  0.77255  0.00000        *
      1   3.07597  4.50235  4.62106  4.08625  3.42277  4.21896  4.65737  2.31224  3.93438  1.10004
3.30196  4.28478  3.73484  4.16061  4.09746  3.56438  3.33416  1.53892  5.22355  4.06250      1 l - - -
          2.68618  4.42225  2.77519  2.73123  3.46354  2.40513  3.72494  3.29354  2.67741  2.69355
4.24690  2.90347  2.73739  3.18146  2.89801  2.37887  2.77519  2.98518  4.58477  3.61503
          0.01786  4.42966  5.15201  0.61958  0.77255  1.02794  0.44277
      2   3.20663  5.35769  3.43699  2.91593  4.74262  3.79684  3.86184  4.19404  2.22051  3.63329
4.51145  3.35190  4.21357  1.03042  1.97661  3.19915  3.40566  3.86211  5.68025  4.45511      2 q - - -
          2.68618  4.42225  2.77519  2.73123  3.46354  2.40513  3.72494  3.29354  2.67741  2.69355
4.24690  2.90347  2.73739  3.18146  2.89801  2.37887  2.77519  2.98518  4.58477  3.61503
          0.01786  4.42966  5.15201  0.61958  0.77255  1.02794  0.44277
      3   3.04778  5.27450  3.21104  2.72757  4.46290  3.69676  1.54312  4.04172  2.28371  3.52674
4.37009  3.20676  4.10842  1.86985  2.41528  3.03486  3.25422  3.70126  5.55932  4.23310      3 h - - -
          2.68618  4.42225  2.77519  2.73123  3.46354  2.40513  3.72494  3.29354  2.67741  2.69355
4.24690  2.90347  2.73739  3.18146  2.89801  2.37887  2.77519  2.98518  4.58477  3.61503
          0.01786  4.42966  5.15201  0.61958  0.77255  1.02794  0.44277
      4   3.58205  5.69643  2.89485  0.43578  5.13702  3.74637  4.43155  4.71884  3.39867  4.25573
5.24565  3.42385  4.38850  3.66070  3.83996  3.52687  3.90322  4.35717  6.19226  4.97107      4 E - - -
          2.68618  4.42225  2.77519  2.73123  3.46354  2.40513  3.72494  3.29354  2.67741  2.69355
4.24690  2.90347  2.73739  3.18146  2.89801  2.37887  2.77519  2.98518  4.58477  3.61503
          0.01786  4.42966  5.15201  0.61958  0.77255  1.02794  0.44277
      5   3.50381  4.81215  5.07087  4.73537  3.82164  4.63206  5.39710  0.57095  4.57956  2.32608
3.70383  4.91178  5.04097  4.89071  4.71907  4.21441  3.80467  2.02234  5.80320  4.57151      5 I - - -
          2.68618  4.42225  2.77519  2.73123  3.46354  2.40513  3.72494  3.29354  2.67741  2.69355
4.24690  2.90347  2.73739  3.18146  2.89801  2.37887  2.77519  2.98518  4.58477  3.61503
          0.01786  4.42966  5.15201  0.61958  0.77255  1.02794  0.44277
      6   3.21492  5.32968  3.30425  2.91546  4.65369  3.75531  3.95311  4.19321  2.38943  3.64875
4.55999  3.36436  4.22972  0.86168  2.41577  3.21879  3.44933  3.86624  5.70361  4.44155      6 q - - -
          2.68618  4.42225  2.77519  2.73123  3.46354  2.40513  3.72494  3.29354  2.67741  2.69355
4.24690  2.90347  2.73739  3.18146  2.89801  2.37887  2.77519  2.98518  4.58477  3.61503
          0.01786  4.42966  5.15201  0.61958  0.77255  0.57743  0.82403
      7   3.27946  5.62872  2.76711  0.82636  5.08526  3.65450  4.20919  4.59605  3.12093  4.11294
4.97087  3.22033  1.96604  3.38024  3.61528  3.22359  3.58365  4.17280  6.21852  4.83300      7 e - - -
          2.68618  4.42225  2.77519  2.73123  3.46354  2.40513  3.72494  3.29354  2.67741  2.69355
4.24690  2.90347  2.73739  3.18146  2.89801  2.37887  2.77519  2.98518  4.58477  3.61503
          0.01462  4.62822  5.35056  0.61958  0.77255  0.80029  0.59638
      8   2.41135  4.99448  3.32252  2.80903  4.26522  1.58782  2.76231  3.67480  2.81203  3.30257
2.55677  2.51859  4.13205  3.16040  3.25541  2.95232  3.13851  3.36279  5.56683  4.25198      8 g - - -
          2.68618  4.42225  2.77519  2.73123  3.46354  2.40513  3.72494  3.29354  2.67741  2.69355
4.24690  2.90347  2.73739  3.18146  2.89801  2.37887  2.77519  2.98518  4.58477  3.61503
          0.01074  4.93473  5.65708  0.61958  0.77255  0.32980  1.26963
      9   3.32265  4.38999  2.93220  2.74585  3.83782  5.04931  3.19681  4.17294  2.47079  3.89219
3.60128  3.31409  3.58016  2.69293  2.82798  1.77681  1.52032  4.31618  5.10809  3.94577      9 t - - -
          2.68618  4.42225  2.77519  2.73123  3.46354  2.40513  3.72494  3.29354  2.67741  2.69355
4.24690  2.90347  2.73739  3.18146  2.89801  2.37887  2.77519  2.98518  4.58477  3.61503
          0.00120  7.12323  7.84557  0.61958  0.77255  0.48576  0.95510
     10   3.31386  3.25661  3.85911  4.11535  2.86280  2.26208  4.45468  2.42647  3.17598  2.36270
3.73679  3.76490  5.81867  3.31890  3.55047  2.70785  3.36862  1.89321  3.78867  2.28909     10 v - - -
          2.68618  4.42225  2.77519  2.73123  3.46354  2.40513  3.72494  3.29354  2.67741  2.69355
4.24690  2.90347  2.73739  3.18146  2.89801  2.37887  2.77519  2.98518  4.58477  3.61503
          0.00120  7.12323  7.84557  0.61958  0.77255  0.48576  0.95510
     11   3.26983  3.45684  2.59324  1.11169  6.08901  5.09817  5.28785  3.25831  3.69222  3.54252
3.33894  3.60964  3.25798  2.49306  4.08269  2.79682  2.93234  2.84428  7.21473  3.05266     11 e - - -
```

```
          2.68618  4.42225  2.77519  2.73123  3.46354  2.40513  3.72494  3.29354  2.67741  2.69355
  4.24690  2.90347  2.73739  3.18146  2.89801  2.37887  2.77519  2.98518  4.58477  3.61503
          0.01401  7.16461  4.33250  0.61958  0.77255  0.48576  0.95510
     12   3.50524  3.73513  4.72777  4.45048  2.13040  2.29346  4.46059  2.02891  3.28241  2.61467
  5.01199  4.14421  5.95318  4.43705  3.20203  4.88070  3.15477  2.50600  4.01685  1.49328     12 y - - -
          2.68618  4.42225  2.77519  2.73123  3.46354  2.40513  3.72494  3.29354  2.67741  2.69355
  4.24690  2.90347  2.73739  3.18146  2.89801  2.37887  2.77519  2.98518  4.58477  3.61503
          0.00116  7.15176  7.87411  0.61958  0.77255  0.75839  0.63190
     13   3.28246  4.86482  3.20323  2.49816  4.71304  3.95655  4.07040  2.97990  2.77651  3.13032
  4.25343  3.45664  5.48448  2.81031  0.92486  2.67387  3.51132  4.42269  7.25495  4.44610     13 r - - -
          2.68618  4.42225  2.77519  2.73123  3.46354  2.40513  3.72494  3.29354  2.67741  2.69355
  4.24690  2.90347  2.73739  3.18146  2.89801  2.37887  2.77519  2.98518  4.58477  3.61503
          0.00114  7.17162  7.89397  0.61958  0.77255  0.34666  1.22773
     14   3.55348  2.61753  6.46774  4.68099  1.29907  3.57891  6.00518  2.62686  4.02557  2.07606
  2.70872  2.60789  6.03521  4.53422  3.95554  4.98289  4.73421  3.09857  3.18212  2.36460     14 f - - -
          2.68618  4.42225  2.77519  2.73123  3.46354  2.40513  3.72494  3.29354  2.67741  2.69355
  4.24690  2.90347  2.73739  3.18146  2.89801  2.37887  2.77519  2.98518  4.58477  3.61503
          0.01522  7.18421  4.24431  0.61958  0.77255  0.48576  0.95510
     15   3.46669  2.76683  6.25783  3.64051  3.82983  5.62954  4.53653  2.65471  3.29987  1.79847
  3.47447  5.72791  5.99518  2.91331  1.72840  3.41611  3.49790  2.82967  4.74025  1.77469     15 r - - -
          2.68618  4.42225  2.77519  2.73123  3.46354  2.40513  3.72494  3.29354  2.67741  2.69355
  4.24690  2.90347  2.73739  3.18146  2.89801  2.37887  2.77519  2.98518  4.58477  3.61503
          0.00114  7.17014  7.89248  0.61958  0.77255  0.33546  1.25530
     16   2.28216  3.67783  4.19142  4.04129  6.12370  3.38730  2.04495  3.82376  2.21296  3.02784
  3.97160  3.69505  4.01014  2.99685  1.56479  2.99571  2.48925  3.28480  7.24069  5.84102     16 r - - -
          2.68618  4.42275  2.77456  2.73174  3.46404  2.40563  3.72545  3.29324  2.67746  2.69255
  4.24740  2.90397  2.73759  3.18197  2.89734  2.37860  2.77538  2.98448  4.58527  3.61553
          0.06219  2.81466  7.90656  0.83152  0.57161  0.48576  0.95510
     17   3.38728  5.19226  3.05411  4.01900  6.18409  4.14541  3.95281  5.67046  0.95111  4.02056
  4.59464  4.15345  2.82921  3.01523  1.62765  2.95795  3.27966  4.45656  7.27080  3.98795     20 k - - -
          2.68618  4.42225  2.77519  2.73123  3.46354  2.40513  3.72494  3.29354  2.67741  2.69355
  4.24690  2.90347  2.73739  3.18146  2.89801  2.37887  2.77519  2.98518  4.58477  3.61503
          0.00113  7.18421  7.90656  0.61958  0.77255  0.48576  0.95510
     18   2.98947  6.84587  0.88318  3.81002  6.19156  2.65888  4.06269  5.67826  4.02809  4.65314
  5.88652  1.62308  5.49879  3.67923  3.89021  2.55321  3.10122  5.23687  7.27806  4.24770     21 d - - -
          2.68618  4.42225  2.77519  2.73123  3.46354  2.40513  3.72494  3.29354  2.67741  2.69355
  4.24690  2.90347  2.73739  3.18146  2.89801  2.37887  2.77519  2.98518  4.58477  3.61503
          0.00113  7.18421  7.90656  0.61958  0.77255  0.48576  0.95510
     19   3.67175  6.95589  3.96321  3.44638  6.29995  0.36546  4.22508  5.78921  3.78767  5.26015
  5.99949  3.72722  5.58305  2.77810  3.49761  3.53364  3.83299  5.34651  7.38628  5.96906     22 G - - -
          2.68618  4.42225  2.77519  2.73123  3.46354  2.40513  3.72494  3.29354  2.67741  2.69355
  4.24690  2.90347  2.73739  3.18146  2.89801  2.37887  2.77519  2.98518  4.58477  3.61503
          0.00604  7.18421  5.24693  0.61958  0.77255  0.48576  0.95510
     20   3.03418  6.83629  2.67642  2.20377  6.18276  2.23034  3.55367  5.66972  2.81568  4.57096
  5.87660  4.56250  5.48909  2.48878  2.92994  1.30062  2.08178  5.22769  7.26813  5.85634     23 s - - -
          2.68618  4.42225  2.77519  2.73123  3.46354  2.40513  3.72494  3.29354  2.67741  2.69355
  4.24690  2.90347  2.73739  3.18146  2.89801  2.37887  2.77519  2.98518  4.58477  3.61503
          0.00113  7.17931  7.90165  0.61958  0.77255  0.41968  1.07077
     21   4.00659  3.87881  5.83501  3.09819  2.74967  4.37608  4.64345  2.40625  3.05993  2.34125
  3.31724  4.43084  2.83997  2.92015  3.05780  3.64288  3.40493  2.80790  2.21386  1.85037     24 y - - -
          2.68618  4.42225  2.77519  2.73123  3.46354  2.40513  3.72494  3.29354  2.67741  2.69355
  4.24690  2.90347  2.73739  3.18146  2.89801  2.37887  2.77519  2.98518  4.58477  3.61503
          0.00113  7.18421  7.90656  0.61958  0.77255  0.48576  0.95510
     22   3.97513  4.71910  6.47747  5.86130  1.88406  4.04912  6.00699  1.64864  3.39495  1.72328
  3.94673  5.83903  6.03646  3.76703  2.49655  4.98462  3.87658  1.62472  6.49744  3.33645     25 v - - -
          2.68619  4.42227  2.77521  2.73125  3.46355  2.40514  3.72496  3.29356  2.67742  2.69356
  4.24632  2.90348  2.73741  3.18148  2.89802  2.37888  2.77507  2.98520  4.58478  3.61505
          0.00345  5.78476  7.90656  0.64622  0.74238  0.48576  0.95510
     23   3.32439  6.45682  3.73045  3.10084  5.66417  3.73361  4.13211  3.48485  3.89782  2.94971
  5.53685  4.79863  2.80142  4.62584  3.47368  2.55650  2.73609  3.00892  0.92351  3.12016     28 w - - -
          2.68618  4.42225  2.77519  2.73123  3.46354  2.40513  3.72494  3.29354  2.67741  2.69355
  4.24690  2.90347  2.73739  3.18146  2.89801  2.37887  2.77519  2.98518  4.58477  3.61503
          0.00113  7.18421  7.90656  0.61958  0.77255  0.48576  0.95510
     24   2.25682  2.51318  6.47665  4.69939  3.81946  3.36854  4.07026  1.87315  5.64295  1.68285
  3.17625  3.56162  6.03635  5.77212  4.21552  4.98447  3.32021  1.39672  6.49751  5.32179     29 v - - -
          2.68618  4.42225  2.77519  2.73123  3.46354  2.40513  3.72494  3.29354  2.67741  2.69355
  4.24690  2.90347  2.73739  3.18146  2.89801  2.37887  2.77519  2.98518  4.58477  3.61503
          0.00113  7.18421  7.90656  0.61958  0.77255  0.48576  0.95510
     25   3.20591  6.77547  2.55295  2.11221  3.98556  5.11772  3.33690  3.46132  3.40635  1.79488
  4.65423  3.03766  5.51050  1.60857  2.32538  4.30886  3.40186  3.61194  4.25462  3.84875     30 q - - -
          2.68618  4.42225  2.77519  2.73123  3.46354  2.40513  3.72494  3.29354  2.67741  2.69355
  4.24690  2.90347  2.73739  3.18146  2.89801  2.37887  2.77519  2.98518  4.58477  3.61503
          0.00113  7.18421  7.90656  0.61958  0.77255  0.48576  0.95510
     26   2.30636  5.88810  3.94377  3.97701  2.15208  5.67509  6.00713  3.03890  5.62169  2.12023
  3.20555  4.66791  6.03907  5.75833  4.52928  2.06107  1.67787  1.92377  4.28996  3.65981     31 t - - -
          2.68618  4.42225  2.77519  2.73123  3.46354  2.40513  3.72494  3.29354  2.67741  2.69355
```

```
4.24690  2.90347  2.73739  3.18146  2.89801  2.37887  2.77519  2.98518  4.58477  3.61503
         0.00112  7.19309  7.91544  0.61958  0.77255  0.48576  0.95510
     27  3.40872  4.99514  2.80171  2.23972  4.83583  3.65289  3.27670  4.53280  2.74867  3.14662
5.88719  2.94308  5.50104  2.08773  2.46346  1.56313  2.92167  3.29389  7.27892  3.38756    32 s - - -
         2.68618  4.42225  2.77519  2.73123  3.46354  2.40513  3.72494  3.29354  2.67741  2.69355
4.24690  2.90347  2.73739  3.18146  2.89801  2.37887  2.77519  2.98518  4.58477  3.61503
         0.00112  7.19309  7.91544  0.61958  0.77255  0.48576  0.95510
     28  1.63323  3.98025  6.45377  3.67443  4.97925  2.60546  6.00903  2.05919  5.62884  2.94701
3.76811  4.18992  6.04037  5.76367  4.11913  2.66743  3.80052  1.50289  2.45142  3.44950    33 v - - -
         2.68618  4.42225  2.77519  2.73123  3.46354  2.40513  3.72494  3.29354  2.67741  2.69355
4.24690  2.90347  2.73739  3.18146  2.89801  2.37887  2.77519  2.98518  4.58477  3.61503
         0.00112  7.19309  7.91544  0.61958  0.77255  0.48576  0.95510
     29  2.47824  4.82813  4.12895  3.07974  3.04007  4.24644  4.16918  3.58647  2.06444  3.85519
4.13730  3.34396  5.51887  2.81487  3.35363  1.76162  2.00314  2.63823  7.23088  2.75455    34 s - - -
         2.68618  4.42225  2.77519  2.73123  3.46354  2.40513  3.72494  3.29354  2.67741  2.69355
4.24690  2.90347  2.73739  3.18146  2.89801  2.37887  2.77519  2.98518  4.58477  3.61503
         0.00112  7.19309  7.91544  0.61958  0.77255  0.48576  0.95510
     30  2.48943  4.22581  5.14031  3.45417  5.40628  3.44420  5.59311  2.79774  3.04598  2.06338
5.34787  5.02391  1.28104  4.29602  2.73211  2.70693  3.18227  2.22250  6.81723  5.56149    35 p - - -
         2.68621  4.42228  2.77522  2.73126  3.46357  2.40516  3.72497  3.29357  2.67744  2.69358
4.24693  2.90350  2.73742  3.18149  2.89804  2.37890  2.77522  2.98460  4.58480  3.61506
         0.02402  4.69434  4.22737  0.25146  1.50358  0.48576  0.95510
     31  2.93792  5.87550  4.53291  3.30758  2.16062  5.66188  3.74469  1.38864  5.60711  2.58627
2.89818  4.52621  6.02589  4.59973  3.54331  4.97182  3.26384  1.41470  4.69825  3.84016    37 i - - -
         2.68605  4.42237  2.77414  2.73135  3.46366  2.40525  3.72506  3.29366  2.67753  2.69367
4.24702  2.90303  2.73751  3.18158  2.89813  2.37899  2.77531  2.98530  4.58489  3.61515
         0.06600  3.93229  3.11736  0.57295  0.82979  0.79290  0.60245
     32  3.69076  4.81267  5.05920  2.24149  3.00290  3.96455  3.63838  2.97524  2.30885  2.91432
3.52086  4.94976  3.15578  3.40706  1.90337  3.09928  2.80255  2.59991  4.71882  2.26055    40 r - - -
         2.68618  4.42225  2.77519  2.73123  3.46354  2.40513  3.72494  3.29354  2.67741  2.69355
4.24690  2.90347  2.73739  3.18146  2.89801  2.37887  2.77519  2.98518  4.58477  3.61503
         0.00818  7.13387  4.91299  0.61958  0.77255  1.38321  0.28871
     33  3.38141  6.62769  1.61417  4.06484  4.51998  3.10317  2.97296  4.47609  4.06274  2.45138
4.21419  1.27676  5.49037  3.87826  2.81557  2.66412  3.85170  4.98194  3.48823  3.91066    41 n - - -
         2.68618  4.42225  2.77519  2.73123  3.46354  2.40513  3.72494  3.29354  2.67741  2.69355
4.24690  2.90347  2.73739  3.18146  2.89801  2.37887  2.77519  2.98518  4.58477  3.61503
         0.00119  7.12688  7.84923  0.61958  0.77255  0.33561  1.25492
     34  3.07906  4.68489  2.70226  1.70457  4.68207  2.67619  5.28222  3.66426  2.90123  3.70194
5.87431  3.47404  1.93424  1.87347  2.96283  2.83949  3.48173  3.73217  7.26612  4.60609    42 e - - -
         2.68672  4.42322  2.77616  2.73220  3.46375  2.40609  3.72591  3.29451  2.67591  2.69416
4.24786  2.90443  2.73836  3.18243  2.89578  2.37958  2.77460  2.98615  4.55432  3.61233
         0.30439  1.40049  4.13828  0.08862  2.46742  0.33200  1.26402
     35  2.59073  6.83354  1.98292  2.69950  3.91962  2.41827  4.10363  4.73945  3.03547  4.44499
4.66720  2.14356  5.48765  2.65353  3.15707  1.88431  2.65641  4.29710  3.95243  3.63408    46 s - - -
         2.68618  4.42225  2.77519  2.73123  3.46354  2.40513  3.72494  3.29354  2.67741  2.69355
4.24690  2.90347  2.73739  3.18146  2.89801  2.37887  2.77519  2.98518  4.58477  3.61503
         0.00849  7.17762  4.86833  0.61958  0.77255  0.56901  0.83491
     36  3.02098  6.83416  2.48918  3.01580  6.18063  1.02736  3.93805  5.66758  2.67433  3.83710
5.87446  2.53624  2.85318  3.18845  3.07513  2.51287  3.82794  4.48270  7.26601  5.85422    47 g - - -
         2.68618  4.42225  2.77519  2.73123  3.46354  2.40513  3.72494  3.29354  2.67741  2.69355
4.24690  2.90347  2.73739  3.18146  2.89801  2.37887  2.77519  2.98518  4.58477  3.61503
         0.04010  7.17715  3.25593  0.61958  0.77255  0.82195  0.57906
     37  2.59070  4.69986  3.63328  1.57595  3.60786  3.08024  3.51586  4.62279  2.10339  3.83585
4.71000  2.82918  5.45372  2.10160  2.31199  3.38557  2.54634  4.08173  7.23203  5.82043    48 e - - -
         2.68618  4.42225  2.77519  2.73123  3.46354  2.40513  3.72494  3.29354  2.67741  2.69355
4.24690  2.90347  2.73739  3.18146  2.89801  2.37887  2.77519  2.98518  4.58477  3.61503
         0.00118  7.13823  7.86058  0.61958  0.77255  0.86735  0.54484
     38  3.14680  6.14058  3.89512  3.08507  3.37086  5.37072  4.61802  2.07270  3.22892  2.41467
4.36470  5.09346  1.93669  3.59019  4.27493  2.70492  2.81647  1.44124  6.71971  4.13587    49 v - - -
         2.68618  4.42225  2.77519  2.73123  3.46354  2.40513  3.72494  3.29354  2.67741  2.69355
4.24690  2.90347  2.73739  3.18146  2.89801  2.37887  2.77519  2.98518  4.58477  3.61503
         0.00116  7.15364  7.87599  0.61958  0.77255  0.21638  1.63697
     39  3.13277  6.82756  2.83115  1.96721  4.26328  5.11279  3.53816  2.70803  3.02340  1.90378
3.44415  3.58435  3.99427  2.36857  2.93325  2.61265  2.73848  2.56981  7.26506  5.85897    50 l - - -
         2.68618  4.42225  2.77519  2.73123  3.46354  2.40513  3.72494  3.29354  2.67741  2.69355
4.24690  2.90347  2.73739  3.18146  2.89801  2.37887  2.77519  2.98518  4.58477  3.61503
         0.00112  7.19309  7.91544  0.61958  0.77255  0.48576  0.95510
     40  3.05683  3.25619  4.27442  2.83390  1.96454  2.10832  3.06175  4.16826  4.19340  2.21731
3.14890  3.10183  5.72057  4.30428  2.85451  3.11605  4.33296  3.27667  6.84141  2.00056    51 f - - -
         2.68618  4.42225  2.77519  2.73123  3.46354  2.40513  3.72494  3.29354  2.67741  2.69355
4.24690  2.90347  2.73739  3.18146  2.89801  2.37887  2.77519  2.98518  4.58477  3.61503
         0.00112  7.19309  7.91544  0.61958  0.77255  0.48576  0.95510
     41  3.56334  2.50029  6.48957  3.29263  1.87208  5.68139  6.01573  1.44907  4.52590  2.03685
3.61939  5.84885  6.04495  5.78270  5.65712  3.49805  3.83547  2.09383  3.70466  2.56498    52 i - - -
         2.68618  4.42225  2.77519  2.73123  3.46354  2.40513  3.72494  3.29354  2.67741  2.69355
4.24690  2.90347  2.73739  3.18146  2.89801  2.37887  2.77519  2.98518  4.58477  3.61503
```

```
          0.00693  7.19309  5.09064  0.61958  0.77255  0.48576  0.95510
     42   3.82526  3.23435  4.73030  4.54271  3.26033  4.47425  4.62623  1.50935  5.63927  1.44688
2.95003  5.83733  6.03796  4.05555  4.05555  3.17985  4.73696  1.38296  6.50112  5.32521     53 v - - -
          2.68618  4.42225  2.77519  2.73123  3.46354  2.40513  3.72494  3.29354  2.67741  2.69355
4.24690  2.90347  2.73739  3.18146  2.89801  2.37887  2.77519  2.98518  4.58477  3.61503
          0.00112  7.18729  7.90963  0.61958  0.77255  0.62231  0.76939
     43   2.06182  1.92975  4.53068  5.76068  3.69553  1.45960  5.98835  3.48068  4.57373  2.86694
4.36344  5.79048  4.52455  4.44140  4.58678  2.46382  4.73334  2.26948  2.25072  5.33305     54 g - - -
          2.68619  4.42274  2.77516  2.73122  3.46298  2.40453  3.72543  3.29315  2.67742  2.69307
4.24739  2.90396  2.73788  3.18116  2.89850  2.37936  2.77464  2.98567  4.58526  3.61431
          0.01006  4.64127  7.90963  2.37681  0.09744  0.62231  0.76939
     44   3.34803  4.59643  6.37685  4.07029  3.51097  5.66096  4.72369  2.01377  4.56050  2.28945
3.23308  3.33031  6.02553  4.17715  3.33783  2.48864  1.98308  1.25514  4.65977  5.33246     70 v - - -
          2.68618  4.42225  2.77519  2.73123  3.46354  2.40513  3.72494  3.29354  2.67741  2.69355
4.24690  2.90347  2.73739  3.18146  2.89801  2.37887  2.77519  2.98518  4.58477  3.61503
          0.00112  7.18729  7.90963  0.61958  0.77255  0.40897  1.09165
     45   3.00343  4.38906  4.01849  3.93761  2.32384  3.33487  2.14329  2.88266  3.28631  2.44745
3.25379  2.19469  5.66617  3.06154  3.65049  2.93901  3.28455  1.90632  6.92554  5.63747     71 v - - -
          2.68618  4.42225  2.77519  2.73123  3.46354  2.40513  3.72494  3.29354  2.67741  2.69355
4.24690  2.90347  2.73739  3.18146  2.89801  2.37887  2.77519  2.98518  4.58477  3.61503
          0.01820  7.19309  4.05779  0.61958  0.77255  0.48576  0.95510
     46   3.44260  6.80692  3.69605  2.67308  3.77996  5.09968  2.62674  3.13028  1.98796  3.07309
2.98279  3.77550  5.49290  2.90188  1.94378  2.48513  2.29149  3.42103  7.24604  2.97083     72 r - - -
          2.68618  4.42225  2.77519  2.73123  3.46354  2.40513  3.72494  3.29354  2.67741  2.69355
4.24690  2.90347  2.73739  3.18146  2.89801  2.37887  2.77519  2.98518  4.58477  3.61503
          0.00114  7.17602  7.89837  0.61958  0.77255  0.31404  1.31115
     47   4.40426  6.44345  0.60708  4.35463  5.64120  5.27171  5.48780  3.78155  3.77032  2.68356
5.52687  2.85252  4.14357  3.84292  3.63966  4.48833  4.63548  1.93945  6.97509  4.55372     73 D - - -
          2.68618  4.42225  2.77519  2.73123  3.46354  2.40513  3.72494  3.29354  2.67741  2.69355
4.24690  2.90347  2.73739  3.18146  2.89801  2.37887  2.77519  2.98518  4.58477  3.61503
          0.00112  7.19309  7.91544  0.61958  0.77255  0.48576  0.95510
     48   3.74330  5.88529  6.48550  5.86954  4.97741  4.67676  3.59727  0.78319  5.65210  1.95445
3.35459  5.84822  3.58645  5.78164  3.11826  4.99438  3.82579  1.73857  6.50779  5.33203     74 i - - -
          2.68618  4.42225  2.77519  2.73123  3.46354  2.40513  3.72494  3.29354  2.67741  2.69355
4.24690  2.90347  2.73739  3.18146  2.89801  2.37887  2.77519  2.98518  4.58477  3.61503
          0.00112  7.19309  7.91544  0.61958  0.77255  0.48576  0.95510
     49   3.30479  4.40745  3.32101  3.32334  6.11459  4.30769  5.31224  5.58440  3.13648  2.50478
3.89534  3.83407  5.51549  4.41661  3.17668  1.51606  0.93826  4.18720  7.23984  5.84344     75 t - - -
          2.68618  4.42225  2.77519  2.73123  3.46354  2.40513  3.72494  3.29354  2.67741  2.69355
4.24690  2.90347  2.73739  3.18146  2.89801  2.37887  2.77519  2.98518  4.58477  3.61503
          0.00112  7.19309  7.91544  0.61958  0.77255  0.48576  0.95510
     50   2.66844  6.84785  1.93362  1.25255  6.19430  3.27689  3.11897  5.68124  3.13959  4.01275
5.88815  3.09380  3.31319  2.26407  3.56817  2.66554  4.56512  3.45265  5.53710  4.00679     76 e - - -
          2.68618  4.42225  2.77519  2.73123  3.46354  2.40513  3.72494  3.29354  2.67741  2.69355
4.24690  2.90347  2.73739  3.18146  2.89801  2.37887  2.77519  2.98518  4.58477  3.61503
          0.00112  7.19309  7.91544  0.61958  0.77255  0.48576  0.95510
     51   2.92848  5.17505  3.30190  2.74311  4.42873  2.87353  2.99738  2.91712  2.72286  2.52998
4.24216  3.26694  4.13789  2.33034  2.33065  1.77181  3.15912  3.51592  5.66569  4.32166     77 s - - -
          2.68618  4.42225  2.77519  2.73123  3.46354  2.40513  3.72494  3.29354  2.67741  2.69355
4.24690  2.90347  2.73739  3.18146  2.89801  2.37887  2.77519  2.98518  4.58477  3.61503
          0.13840  5.08763  2.09493  0.61958  0.77255  0.48576  0.95510
     52   2.86071  5.34660  3.09603  1.98714  4.69491  2.82721  2.36813  4.16978  1.85538  3.64451
4.39065  3.09063  2.88864  2.89780  2.49258  2.82421  3.08889  3.73993  5.76901  4.37480     78 k - - -
          2.68618  4.42225  2.77519  2.73123  3.46354  2.40513  3.72494  3.29354  2.67741  2.69355
4.24690  2.90347  2.73739  3.18146  2.89801  2.37887  2.77519  2.98518  4.58477  3.61503
          0.17272  4.83261  1.89270  0.61958  0.77255  0.80029  0.59638
     53   2.28197  4.92690  3.33170  2.75037  4.14265  3.67933  3.84525  3.54822  2.56269  2.73319
4.00646  3.23902  4.06144  2.14666  1.76869  2.90600  3.06662  3.25569  3.23086  4.12019     79 r - - -
          2.68618  4.42225  2.77519  2.73123  3.46354  2.40513  3.72494  3.29354  2.67741  2.69355
4.24690  2.90347  2.73739  3.18146  2.89801  2.37887  2.77519  2.98518  4.58477  3.61503
          0.01397  4.67385  5.39620  0.61958  0.77255  1.02794  0.44277
     54   2.37372  5.43527  0.77418  2.48361  5.00731  3.43345  4.15192  4.45136  3.22790  4.06892
4.96189  3.04055  4.10765  3.34440  3.81124  3.06451  3.46845  4.01132  6.24457  4.78688     80 d - - -
          2.68618  4.42225  2.77519  2.73123  3.46354  2.40513  3.72494  3.29354  2.67741  2.69355
4.24690  2.90347  2.73739  3.18146  2.89801  2.37887  2.77519  2.98518  4.58477  3.61503
          0.01786  4.42966  5.15201  0.61958  0.77255  1.02794  0.44277
     55   2.54526  4.51060  3.59867  3.37633  4.69947  3.22950  4.51898  4.19407  3.48575  3.84907
4.67356  3.56363  2.63620  3.78347  3.80186  0.73002  3.04298  3.58978  6.01087  4.77962     81 s - - -
          2.68618  4.42225  2.77519  2.73123  3.46354  2.40513  3.72494  3.29354  2.67741  2.69355
4.24690  2.90347  2.73739  3.18146  2.89801  2.37887  2.77519  2.98518  4.58477  3.61503
          0.01786  4.42966  5.15201  0.61958  0.77255  1.02794  0.44277
     56   0.44114  4.57798  4.13902  3.99945  4.71762  3.38090  4.93763  3.93826  4.03369  3.78910
4.78232  3.97307  4.17446  4.32383  4.21642  2.93621  3.25373  3.48461  6.07941  4.96190     82 A - - -
          2.68618  4.42225  2.77519  2.73123  3.46354  2.40513  3.72494  3.29354  2.67741  2.69355
4.24690  2.90347  2.73739  3.18146  2.89801  2.37887  2.77519  2.98518  4.58477  3.61503
          0.01786  4.42966  5.15201  0.61958  0.77255  1.02794  0.44277
```

```
     57   2.58927  4.50441  3.73450  3.45533  4.50433  3.30041  4.50546  3.83358  3.45383  3.57200
4.46349  3.63051  2.66037  3.79154  3.74290  2.77286  0.78768  3.37607  5.86255  4.65997     83 t - - -
          2.68618  4.42225  2.77519  2.73123  3.46354  2.40513  3.72494  3.29354  2.67741  2.69355
4.24690  2.90347  2.73739  3.18146  2.89801  2.37887  2.77519  2.98518  4.58477  3.61503
          0.01786  4.42966  5.15201  0.61958  0.77255  1.02794  0.44277
     58   3.76402  5.49036  4.32136  3.75546  4.99690  4.09465  4.39898  4.61287  2.68382  4.02573
5.03930  4.05717  4.59601  3.61696  0.38602  3.84020  4.00824  4.31842  5.86623  4.83751     84 R - - -
          2.68618  4.42225  2.77519  2.73123  3.46354  2.40513  3.72494  3.29354  2.67741  2.69355
4.24690  2.90347  2.73739  3.18146  2.89801  2.37887  2.77519  2.98518  4.58477  3.61503
          0.01786  4.42966  5.15201  0.61958  0.77255  1.02794  0.44277
     59   1.50036  4.37769  3.97205  3.66486  4.49042  3.22700  4.61392  3.68029  3.63865  3.53264
4.42281  3.70214  3.99020  3.92808  3.89763  2.68841  1.01115  3.22459  5.89600  4.71406     85 t - - -
          2.68618  4.42225  2.77519  2.73123  3.46354  2.40513  3.72494  3.29354  2.67741  2.69355
4.24690  2.90347  2.73739  3.18146  2.89801  2.37887  2.77519  2.98518  4.58477  3.61503
          0.01786  4.42966  5.15201  0.61958  0.77255  1.02794  0.44277
     60   2.96551  3.54295  3.56377  2.92676  4.22317  3.72407  2.88271  3.70182  2.31891  3.27555
4.14293  3.34550  4.12292  3.01262  1.13184  3.03253  3.18724  3.40555  5.40808  4.13872     86 r - - -
          2.68618  4.42225  2.77519  2.73123  3.46354  2.40513  3.72494  3.29354  2.67741  2.69355
4.24690  2.90347  2.73739  3.18146  2.89801  2.37887  2.77519  2.98518  4.58477  3.61503
          0.01786  4.42966  5.15201  0.61958  0.77255  1.02794  0.44277
     61   3.85085  5.16685  5.08705  4.76253  3.45479  4.68770  5.25265  2.71567  4.53131  0.40420
3.36014  4.98987  5.04817  4.75685  4.60220  4.38129  4.13003  2.89097  5.49731  4.33900     87 L - - -
          2.68618  4.42225  2.77519  2.73123  3.46354  2.40513  3.72494  3.29354  2.67741  2.69355
4.24690  2.90347  2.73739  3.18146  2.89801  2.37887  2.77519  2.98518  4.58477  3.61503
          0.01786  4.42966  5.15201  0.61958  0.77255  1.02794  0.44277
     62   3.85085  5.16685  5.08705  4.76253  3.45479  4.68770  5.25265  2.71567  4.53131  0.40420
3.36014  4.98987  5.04817  4.75685  4.60220  4.38129  4.13003  2.89097  5.49731  4.33900     88 L - - -
          2.68618  4.42225  2.77519  2.73123  3.46354  2.40513  3.72494  3.29354  2.67741  2.69355
4.24690  2.90347  2.73739  3.18146  2.89801  2.37887  2.77519  2.98518  4.58477  3.61503
          0.01206  4.42386        *  0.61958  0.77255  0.00000        *
//
```
